# Functional connectivity across the human subcortical auditory system using an autoregressive matrix-Gaussian copula graphical model approach with partial correlations

**DOI:** 10.1101/2022.09.15.508099

**Authors:** Noirrit Kiran Chandra, Kevin R. Sitek, Bharath Chandrasekaran, Abhra Sarkar

## Abstract

The auditory system comprises multiple subcortical brain structures that process and refine incoming acoustic signals along the primary auditory pathway. Due to technical limitations of imaging small structures deep inside the brain, most of our knowledge of the subcortical auditory system is based on research in animal models using invasive methodologies. Advances in ultra-high field functional magnetic resonance imaging (fMRI) acquisition have enabled novel non-invasive investigations of the human auditory subcortex, including fundamental features of auditory representation such as tonotopy and periodotopy. However, functional connectivity across subcortical networks is still underexplored in humans, with ongoing development of related methods. Traditionally, functional connectivity is estimated from fMRI data with full correlation matrices. However, partial correlations reveal the relationship between two regions after removing the effects of all other regions, reflecting more direct connectivity. Partial correlation analysis is particularly promising in the ascending auditory system, where sensory information is passed in an obligatory manner, from nucleus to nucleus up the primary auditory pathway, providing redundant but also increasingly abstract representations of auditory stimuli. While most existing methods for learning conditional dependency structures based on partial correlations assume independently and identically Gaussian distributed data, fMRI data exhibit significant deviations from Gaussianity as well as high temporal autocorrelation. In this paper, we developed an autoregressive matrix-Gaussian copula graphical model (ARMGCGM) approach to estimate the partial correlations and thereby infer the functional connectivity patterns within the auditory system while appropriately accounting for autocorrelations between successive fMRI scans. Our results show strong positive partial correlations between successive structures in the primary auditory pathway on each side (left and right), including between auditory midbrain and thalamus, and between primary and associative auditory cortex. These results are highly stable when splitting the data in halves according to the acquisition schemes and computing partial correlations separately for each half of the data, as well as across cross-validation folds. In contrast, full correlation-based analysis identified a rich network of interconnectivity that was not specific to adjacent nodes along the pathway. Overall, our results demonstrate that unique functional connectivity patterns along the auditory pathway are recoverable using novel connectivity approaches and that our connectivity methods are reliable across multiple acquisitions.

## 1. Introduction

The mammalian auditory pathway conveys acoustic information between the inner ear—where sounds are mechanoelectrically transduced in the cochlea—and auditory cortex by way of multiple subcortical nuclei across the brainstem, midbrain, and thalamus. Much of our knowledge of the auditory system arises from anatomical and physiological research with non-human animal models (McIntosh and Gonzalez-Lima, 1991; Popper and Fay, 1992; Webster *et al*., 1992). This work has contributed tremendously to our understanding of the mammalian auditory system. However, due to methodological and ethical limitations, our ability to directly assess auditory function in the human nervous system is severely constrained (as discussed in (Moerel *et al*., 2021).

This is particularly true for subcortical auditory structures. In mammals, the ascending central auditory pathway receives signals from the cochlea of the inner ear by way of the cochlear nerve, which principally innervates the cochlear nucleus in the brainstem. Auditory signals are then transmitted to the superior olivary complex, which is the first decussation point at which signals largely pass contralaterally from the left cochlear nucleus to the right superior olive (and similarly from right to left). From the superior olive, auditory signals travel (via the lateral lemniscus) to the inferior colliculus in the midbrain. The last subcortical auditory structure is the medial geniculate nucleus of the thalamus, which then passes information to primary auditory cortex. In addition to the ascending “lemniscal” auditory pathway, an equal number of efferent connections transmit top-down information from higher order auditory regions to earlier auditory structures (Malmierca and Ryugo, 2011; Winer, 2005). Due to the small size of the subcortical auditory structures—tightly packed in with other heterogeneous nuclei and white matter pathways—and their anatomical location deep within the cranium, the subcortical auditory structures have received limited attention in non-invasive human research.

Functional magnetic resonance imaging (fMRI) is the most popular non-invasive method for probing macroscopic network-related brain activity. While studies of the human subcortical auditory system are somewhat limited, previous task-based fMRI research has functionally localized the subcortical auditory structures (Sitek *et al*., 2019), identified the tonotopic frequency mappings within the auditory midbrain and thalamus (De Martino *et al*., 2013; Moerel *et al*., 2015; Ress and Chandrasekaran, 2013), separated top-down and bottom-up speech-selective subregions of auditory thalamus (Mihai *et al*., 2019; Tabas *et al*., 2021), and recorded level-dependent BOLD signals throughout the auditory pathway (Hawley *et al*., 2005; Sigalovsky and Melcher, 2006).

In contrast to task-related BOLD activity, fMRI connectivity methods (often utilizing resting state fMRI paradigms and full correlation analysis) are commonly used to assay cortical brain networks (Biswal *et al*., 1995; Gordon, Laumann, Adeyemo, *et al*., 2017; Power *et al*., 2011; Smith, Beckmann, *et al*., 2013), including the cortical auditory system (Abrams *et al*., 2020; Cha *et al*., 2016; Chen *et al*., 2020; Eckert *et al*., 2008; Maudoux *et al*., 2012; Ren *et al*., 2021). However, fMRI connectivity methods have limited history in subcortical research, especially in the auditory system, where they have—to our knowledge—only been utilized a handful of times to assess connectivity differences between individuals with and without tinnitus percepts (Berlot *et al*., 2020; Hofmeier *et al*., 2018; Leaver *et al*., 2016; Zhang *et al*., 2015).

From the seminal resting state connectivity studies identifying human default mode and motor networks (Biswal *et al*., 1995; Fox and Raichle, 2007), to work linking functional connectivity with behavioral variability (Baldassarre *et al*., 2012; Deng *et al*., 2016), to investigations into brain network differences associated with disorders (Chai *et al*., 2016; Di Martino *et al*., 2011; Ferri *et al*., 2018; Greicius *et al*., 2007; Hahn *et al*., 2011; Husain and Schmidt, 2014; Kaiser *et al*., 2015; Sitek *et al*., 2016; Wilson *et al*., 2022), full correlation analysis has contributed tremendously to our understanding of human brain networks. However, moving beyond the traditional full correlation analysis should enable greater specificity in assessing functional connectivity patterns, particularly in identifying specific node-to-node connectivity patterns within an established brain network (Marrelec *et al*., 2006; Smith, 2012). In contrast to full correlations, which represent both direct and indirect connections, partial correlation analyses represent the direct association between two specific nodes after filtering out the effects of the remaining nodes and thus hold great promise for estimating direct functional connectivity within a network (Smith, Beckmann, *et al*., 2013). (Please refer to the illustration in the Supplementary Materials.) For these reasons, partial correlation approaches are increasingly used to study functional connectivity networks in the brain (Wang *et al*., 2016; Warnick *et al*., 2018), including improved identification of individualized connectivity profiles compared to full correlation methods (Menon and Krishnamurthy, 2019), as well as improved prediction of brain disorders (Reeves *et al*., 2023; Skåtun *et al*., 2017; de Vos *et al*., 2018) and identification of general cognitive ability (Sripada *et al*., 2021). However, we are unaware of the application of such methods to assess functional connectivity within subcortical networks, particularly within the human auditory system.

In this article, we build on the Bayesian precision factor model (PFM) introduced recently in Chandra et al. (2021) to develop a novel highly robust autoregressive matrix-Gaussian copula graphical model (ARMGCGM) to assess partial correlation-based functional connectivity in a specific network in the human brain that spans subcortical and cortical regions, the auditory system. The PFM decomposes the model precision matrix into a flexible low-rank and diagonal structure, then exploits that to design very efficient estimation algorithms. However, it makes the restrictive assumption that each variable is marginally Gaussian distributed. Several studies in the literature also make this assumption (Yu *et al*., 2022; Zhang *et al*., 2014) which has the very useful implication that a zero partial correlation between two variables (equivalent to a zero entry in the precision matrix in the corresponding position) also means independence between them after removing the effects of other variables. However, in many applications—including ours—data are often not Gaussian distributed. Additionally, data from successive fMRI volumes exhibit strong autocorrelation. The ARMGCGM extends the PFM to the case where the univariate marginals can be any arbitrary distribution while also accounting for the autocorrelations between successive fMRI scans. The association between the variables are modeled using a Gaussian copula that implies conditional independence for zero partial correlation, allowing easy interpretability of the conditional dependence graph. As developed, the ARMGCGM approach is broadly applicable for studying undirected functional graphs using large-scale fMRI data.

We use the novel ARMGCGM to investigate functional connectivity between specific nodes across the human auditory system. We used publicly available 7T resting state fMRI from over one hundred participants to examine auditory connectivity. To probe connectivity within the auditory system, we included auditory cortical regions of interest as well as subcortical auditory regions derived from human histology (Sitek *et al*., 2019). As the auditory pathway comprises a chain of multiple subcortical structures, and due to the largely lateralized organization of the lemniscal auditory system, we hypothesized that partial correlations would be greatest between adjacent nodes in the same hemisphere, over and above the contributions from other auditory (and non-auditory) regions of interest. In particular, because of the critical position of auditory midbrain and thalamus as computational hubs involving bottom-up and top-down information transfer (Mihai *et al*., 2019; Tabas *et al*., 2021), we hypothesized strong partial correlations between inferior colliculus and medial geniculate. We further assessed reliability across acquisitions by separately analyzing data with anterior–posterior and posterior–anterior phase-encoding directions, as well as leave-10%-out cross-validation. We then compared our ARMGCGM-based connectivity results with those from a full correlation approach as well as alternative partial correlation methods. Overall, our consistent findings of hierarchical connectivity within the auditory system using our novel partial correlation method—consistent across data partitions and leave-10%-out validation— demonstrate the methodological reliability of our ARMGCGM approach as well as the neurobiological organization of auditory structures in the human primary auditory system.

## 2. Materials and methods

### 2.1 Magnetic resonance imaging acquisition and processing

We used resting state fMRI from 106 participants in the 7T Human Connectome Project (Elam *et al*., 2021; Van Essen *et al*., 2012). Specifically, we utilized the minimally preprocessed volumetric data in common space (Glasser *et al*., 2013). BOLD fMRI data were acquired with 1.6 mm isotropic voxel size across four runs of a resting state paradigm (repetition time [TR] = 1 s, 900 TRs per run). Two runs were acquired with anterior–posterior (AP) phase encoding, and two were acquired with posterior–anterior (PA) phase encoding. For all ROIs and runs, we discarded the first 50 TRs to increase stability.

For each individual and each run, we extracted mean timeseries from predefined regions of interest (ROIs). Subcortical auditory ROIs were defined using the Sitek–Gulban atlas (Sitek *et al*., 2019). Cortical ROIs were defined using FreeSurfer’s implementation of the DKT atlas (Dale *et al*., 1999; Klein and Tourville, 2012). For this study we used transverse temporal gyrus (TTG) and superior temporal gyrus (STG) as auditory ROIs, as well as pericalcarine cortex (Calc) and superior frontal gyrus (SFG) as non-auditory control ROIs (see Supplementary Materials). Mean timeseries were extracted for each ROI using nilearn’s [Masker] function.

### 2.2 Data partitioning and cross-validation

BOLD fMRI is prone to geometric distortions in the phase-encoding direction which can be largely corrected using a variety of methods (Esteban *et al*., 2021; Jezzard and Balaban, 1995). Adjacent to motion-sensitive cerebrospinal fluid (CSF), the brainstem is particularly susceptible to such geometric distortions. Although the HCP minimal preprocessing pipeline corrects for phase-encoding distortions (Glasser *et al*., 2013), to isolate the potential residual contribution of phase encoding direction on functional connectivity estimates, we conducted separate analyses on data collected with posterior–anterior (PA) phase encoding direction (runs 1 and 3) and anterior– posterior (AP) phase encoding direction (runs 2 and 4) and compared the results. As the fMRI data acquired in the two phases will be analyzed separately but using the same probability-model, we use the same notations for the different phases to describe our proposed model in the following sections.

We further evaluated our results using a leave-10%-out cross-validation (CV) approach. We assessed the stability of the connectivity estimates by checking the correlation between the results obtained from each acquisition scheme.

### 2.3 Autoregressive matrix-Gaussian copula graphical models

The Bayesian precision factor model (PFM), developed recently in (Chandra *et al*., 2021), provided a novel computationally efficient robust technique for estimating precision matrices. Since partial correlations can be readily obtained from the precision matrix, the approach allowed straightforward estimation of the underlying connectivity graphs. Previously, (Lee and Kim, 2021) obtained the precision matrix by inverting the estimated covariance matrix. However, this approach often tends to exhibit poor empirical performance (Pourahmadi, 2013).

Most partial correlation-based conditional dependency graph estimation procedures in the statistical literature (Cai *et al*., 2020; Chandra *et al*., 2021; Friedman *et al*., 2008; Warnick *et al*., 2018) assume that the joint distribution of the data is independent and identically distributed (iid) multivariate Gaussian, which implies that the univariate marginal distributions are also all Gaussians. However, successive fMRI scans have strong autocorrelations, and their marginal distributions exhibit substantial deviance from the Gaussian assumption. In this paper, we therefore extend the PFM to accommodate non-Gaussian marginals, while appropriately accounting for the temporal dependence between successive fMRI scans.

Let 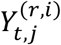 be the fMRI timeseries corresponding to the *i*-th individual’s *j*-th ROI at the *t*-th timepoint in the *r*-th run. In our application we are interested in studying the connectivity between *d* = 12 ROIs along the central auditory pathway using fMRI timeseries of length *T* = 850 from *N* = 106 individuals each undergoing *R* = 2 runs. We let 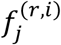 be the (unknown) marginal density of 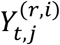 with corresponding cumulative distribution function (CDF) 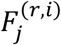.

Copulas provide a broadly applicable class of tools that allow the joint distribution of 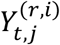 to be flexibly characterized by first modeling the univariate marginals 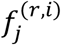 and then hierarchically modeling their joint dependencies by mapping the 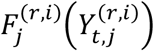’s to a joint probability space. For Gaussian copulas, this is done by setting 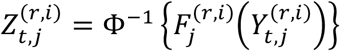 where Φ(·) is the CDF of a standard Gaussian distribution. This implies marginally 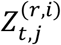 ∼ N(0,1) for all *r, i, j, t*, where N(*μ, σ*^2^) denotes a univariate Gaussian distribution with mean *μ* and variance *σ*^2^. We let 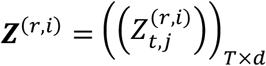 denote the matrix of fMRI signals corresponding to the *i*-th individual in the *r*-th run in the transformed Gaussian space. The Gaussian copula assumption on 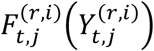’s implies that the joint distribution of ***Z***^(*r,i*)^ is Gaussian as well. The dependencies between the 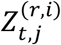’s are therefore characterized entirely by their correlations. Additionally, since the dependence relationships between the observed 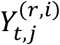’s are modeled only through 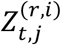’s, these correlations also completely characterize the dependencies between the 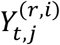’s.

Note that probabilistic dependencies exist between 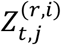’s across both *j* and *t*; dependence across *j* incurs due to the interaction between the ROIs whereas dependence across *t* occurs due to the temporal dependence between successive fMRI scans. We let ***R***_***Ω***_ denote the *d × d* correlation matrix accounting for the dependence across the *d* different ROIs across all runs and individuals.

While our main interest lies in estimating these dependencies between the ROIs, it is also crucial to consider the temporal dependence in the 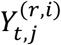’s. Let 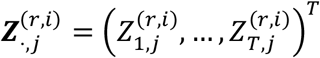 be the *j*-th column of ***Z***^(*r,i*)^, i.e., the timeseries corresponding to the *i*-th individual’s *j*-th ROI in the *r*-th run. We develop our model in a hierarchical manner. To being with, we use autoregressive (AR) processes of order *L*to model higher-order temporal dependencies in the 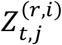’s as

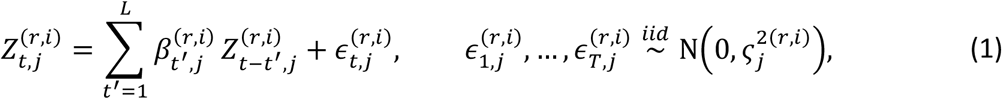

with 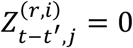 if *t’* ≥ *t*. We assume separate (***β, ς***^2^) parameters across (*r, i*, *j*) in (1) to make the model adapt to different timeseries patterns across different ROIs, individuals, and runs. Let 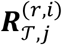 be the correlation matrix of 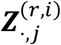 induced by the AR(*L*) model in (1). We let 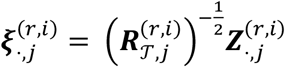 be the AR corrected timeseries corresponding to the *j*-th ROI for *j* = 1, ..., *d*. We define the *T* × *d* matrix 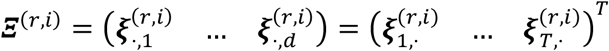 where 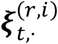 is the *t*-th row of **Ξ** ^(***r, i***)^ and can be interpreted as the fMRI signals in the Gaussian copula space subsequent to filtering out the temporal dependence at time point *t*. We then let

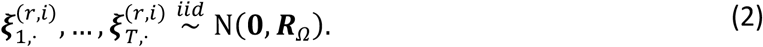

The formulations in (1)-(2) imply the following joint distribution on ***Z***^(*r, i)*^

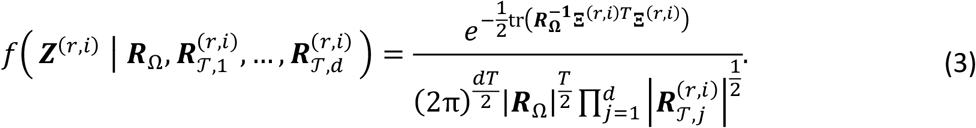

Although the 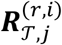’s are massive dimensional *T* × *T* matrices, notably they are characterized entirely by the associated autoregressive parameters 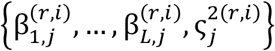, leading to their straightforward numerically inexpensive evaluations in (3); we discuss the details in the posterior computation section in Supplementary Materials. Let **Ω *=*** ((ω_*j, j* ’_)), be the precision matrix corresponding to ***R***_Ω,_ i.e.,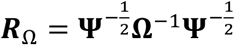 with **Ψ** = diag(**Ω**^-1^)Then, by properties of copula and the simple multivariate Gaussian distribution, it can be shown that, irrespective of the form of the marginal 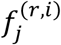’s, the ROIs 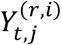 and 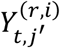 will be conditionally dependent on (and hence functionally connected with) each other given the rest if and only if 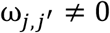. This way, the precision matrix **Ω** characterizes the functional connectivity between the different 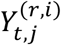’s across *j*.

We model the unknown univariate marginal distributions 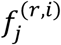’s using location-scale mixtures of Gaussians. Such mixtures can flexibly estimate a very large class of unknown densities (Ghosal *et al*., 1999). Specifically, we let

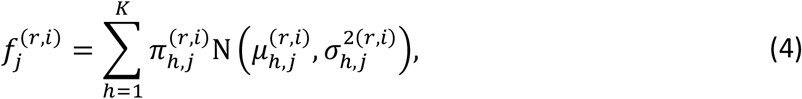

where 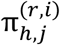 is the weight attached to the *h*-th mixture component and 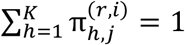 with *K* being a suitably chosen, moderately large, fixed integer.

While fMRI timeseries often showcase very distinct patterns and distributions for different individuals as well as across different ROIs of the same individual, we expect primary sensory region to be highly similar between healthy participants due to strongly stereotyped processing across individuals and well-conserved auditory function in these brain regions across evolution (Hutchison *et al*., 2013). In (4), we therefore consider separate mixture models across different ROIs, individuals, and runs, whereas in (3), the ***Z***^(*r, i*)^’s share a common correlation matrix ***R***_Ω_ across all individuals and runs. This modeling strategy also allows borrowing of information across individuals and runs to amplify signals for estimating resting-state functional connectivity in adult humans while accounting for subject and run-specific variabilities using separate marginal distributions. This is conceptually similar to approaches used in group independent component analysis (Calhoun *et al*., 2001) and cohort-level brain mapping (Varoquaux *et al*., 2013). In later sections, we showed that this approach yields highly consistent estimates of functional connectivity graphs even though BOLD signals from small deep brain regions have very low signal-to-noise ratio (Bianciardi *et al*., 2016; Colizoli *et al*., 2020; de Hollander *et al*., 2017; Sclocco *et al*., 2018).

To estimate the precision matrix **Ω**, we consider the framework of Chandra et al. (2021). Recall that 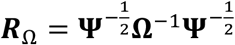 with **Ψ** = diag(Ω^-1^). We assume **Ω** to admit a lower-rank plus diagonal (LRD) decomposition **Ω = ΛΛ^*T*^ + + Δ** where **Λ** is a *d* × *q* matrix and a diagonal matrix 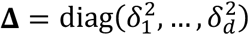 with positive 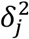’s. Notably, all positive definite matrices admit such a representation for some *q* ≤ *d*.

We take a Bayesian route to estimation and inference, where we assign priors to the model parameters, and then infer them based on samples drawn from the posterior using a Markov chain Monte Carlo (MCMC) algorithm discussed in detail in the Supplementary Materials. As was shown in Chandra et al. (2021), the LRD representation makes the MCMC sampling very efficient via a latent variable augmentation scheme. In the Supplementary Materials, we also discuss a multiple hypothesis testing-based edge discovery procedure that utilizes the posterior uncertainty of the parameters and controls the false discovery rate (FDR) at 10% level in a principled manner.

### 2.4 Priors on the parameters

For the autoregressive parameters in equation (*1*), for all *r, i, j*,we assume 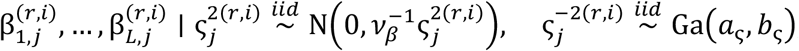 where 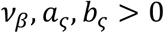 are fixed hyperparameters, and Ga(*a, b*) denotes a gamma distribution with mean *a*/*b* and variance *a*/*b*^2^. For the parameters of the mixture models specifying the marginals in equation (4), we consider the following priors

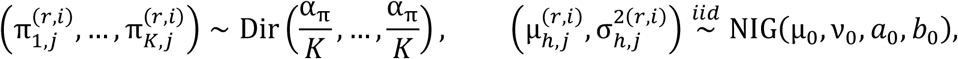

where (*μ, σ*^***2***^) ∼ **NIG** (*μ_0_, ν_0_, *a*_0_, *b*_0_*), implying that 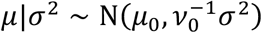 and *σ*^*-****2***^∼ Ga(*a*_0_, *b*_0_).

We consider a shrinkage prior on the elements of **Λ** that shrinks redundant elements of **Λ** to zero allowing additional model-based parameter reduction. In particular, we assign a two-parameter generalization of the Dirichlet-Laplace (DL) prior from Bhattacharya et al. (Bhattacharya et al., 2015) that allows more flexible tail behavior on the elements of **Λ**. On a *d*-dimensional vector ***θ*** = (θ …,…, θ_*d*_)^*T*^, our DL prior with parameters *a* and *b*, denoted by DL(*a, b*), can be specified in the following hierarchical manner manner

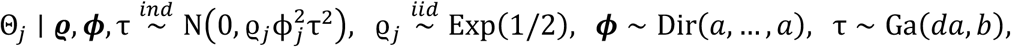

where *θ*_*j*_ is the *j*-th element of ***θ, ϕ*** and ***ϱ*** are vectors of same length as ***θ***, Exp(*a*) is an exponential distribution with mean 1/*a*, Dir(*a*_1_,…, *a*_d_) is a *d*-dimensional Dirichlet distribution. The original DL prior is a special case with *b* = 1/2. We let vec(**Λ**) ∼ DL(*a, b*).

We use a Dirichlet process (DP) prior (Ferguson, 1973) on the 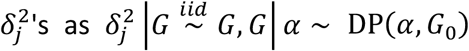 With ***G***_0_ ***=*** Ga(*a*_δ_, *b*_*δ*_), *α* ∼ Ga(*a*_α_*b*_*α*_), where *α* is the concentration parameter and ***G***_0_ is the base measure to favor a smaller number of unique 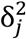’s in a model-based manner. The DP model allows clustering the 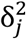’s facilitating additional data-adaptive parameter reduction when necessary. Additionally a DP prior has full prior support on the range-space of the number of unique 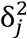’s implying a fully flexible model (see Chapter 4 of (Ghosal and van der Vaart, 2017).

We discuss a Markov chain monte Carlo (MCMC) based strategy of sampling from the posterior of ARMGCGM in Section S1.2 of the Supplementary Materials, where the involving steps are parallelized over the subjects, allowing scalability. In Section S1.3 of the Supplementary Materials, we discuss the choice of hyperparameters used for the analyses presented in this paper. Our implementation using the proposed parallelized MCMC scheme ran in 125 min in a system with 13th Gen Intel(R) Core(TM) i9-13900K CPU and 128GB RAM, fitting the ARMGCGM to the 12-node auditory network with 7,500 MCMC iterations, *including* both phase-encoding schemes.

## 3. Results

### 3.1 Partial correlations between regions of interest

Using our ARMGCGM approach, we first estimate the precision matrix ***R*** and subsequently compute the partial correlation matrix. We report the significant edges subject to controlling the posterior false discovery rate (Sarkar *et al*., 2008) at the 10% level. (Details are provided in Section S1.4 in the Supplementary Materials.) In Figure 2 we provide the circos plots of the connectivity graphs along with respective weighted adjacency matrices. The (*j,h*)-th off-diagonal element of the adjacency matrices admit the value 0 if the edge between ROIs *j* and *h* is not statistically significant, if the edge is significant then the corresponding partial correlation is plugged in to indicate the strength of the edge.

**Figure 1.**
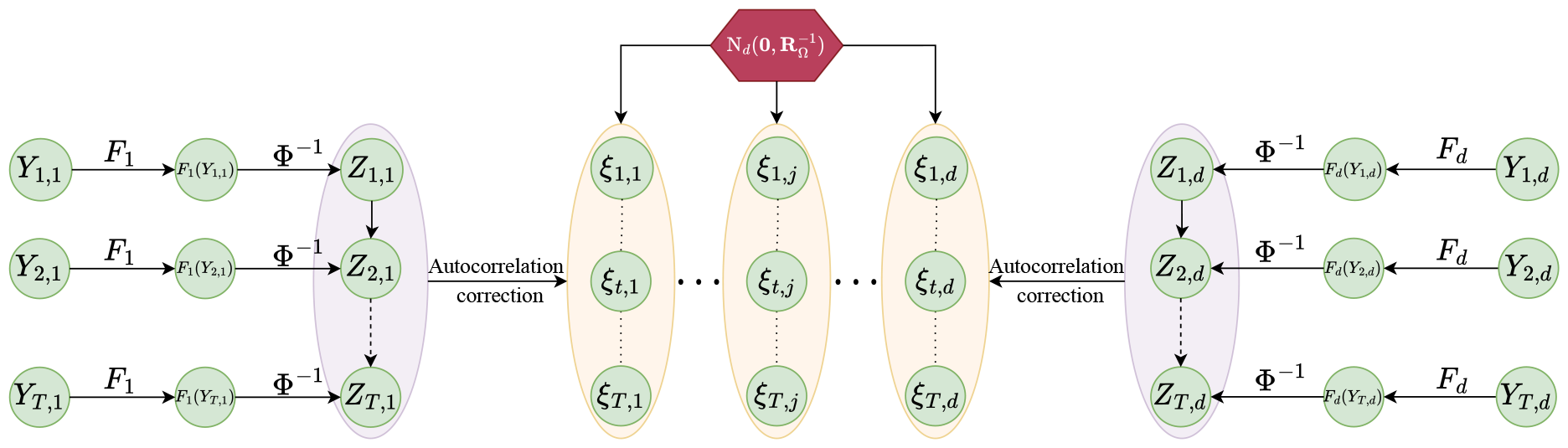
Graphical illustration of the hierarchical structure of the autoregressive matrix-Gaussian copula graphical model (ARMGCGM): For brevity, we omit the superscript (r,i)—the run and subject indicators, respectively, in this illustration. Y_t, j_ is the observed BOLD signal from the j-th ROI at the t-th time point. Z_t,j_ = Φ^−1^ (F_j_(Y_t,j_)) is the BOLD signal in the Gaussian copula space. Probabilistic dependencies exist between Z_t,j_’s across both j and t; dependence across j incurs due to the interaction between the ROIs whereas dependence across t occurs due to the temporal dependence between successive fMRI scans. ξ_t,j_’s are the autocorrelation corrected Z_t,j_’s with 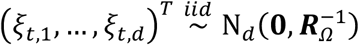for all t=1,...,T.

**Figure 2.**
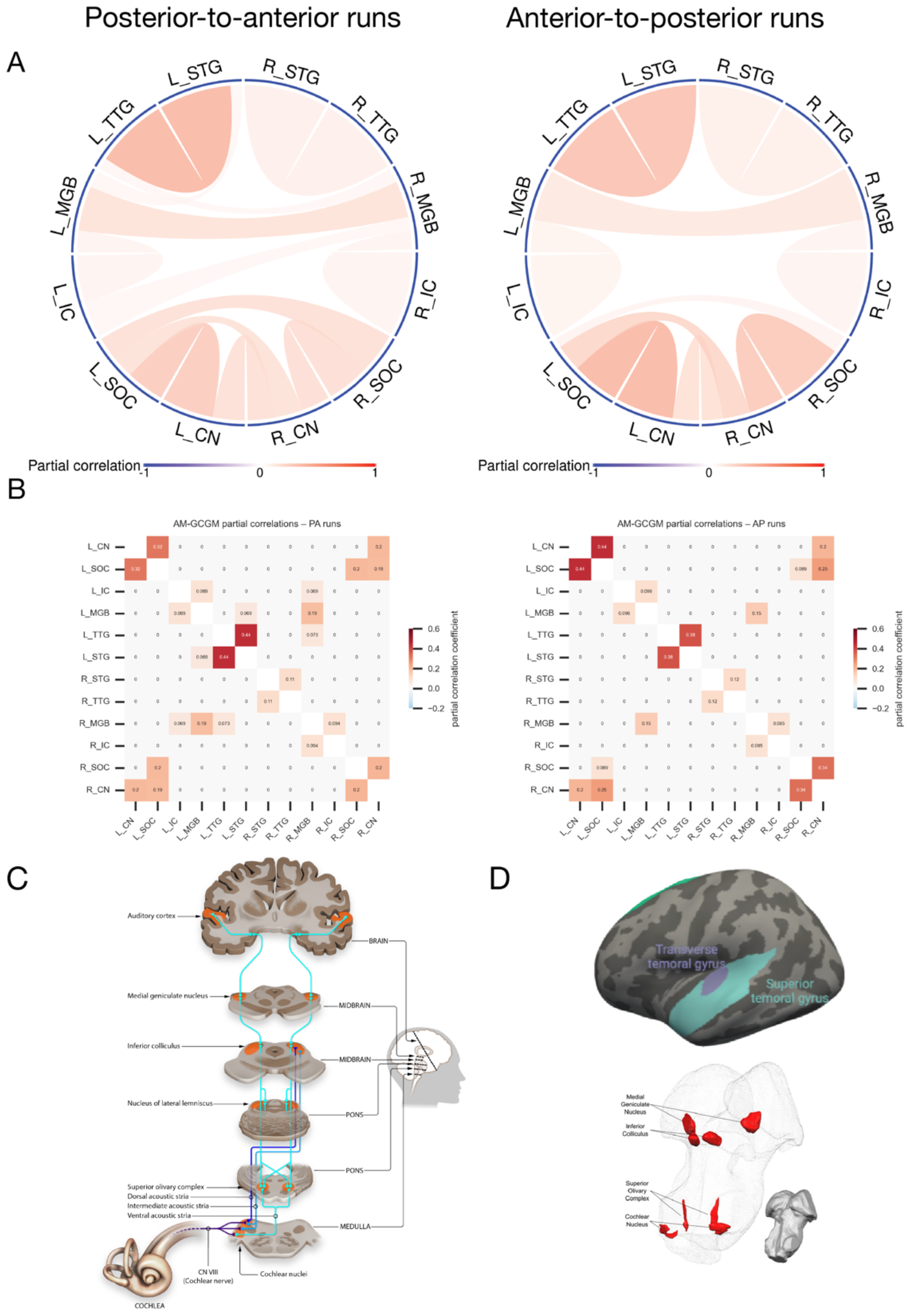
A: Partial correlation connectivity in data acquired with posterior-to-anterior (PA; left) and anterior-to-posterior (AP; right) phase-encoding directions using the ARMGCGM approach in subcortical and cortical auditory regions. Positive (negative) associations are represented by red (blue) links, their opacities being proportional to the corresponding association strengths. The link widths are inversely proportional to the number of edges associated with the corresponding nodes. B: The same results as (A), viewed as adjacency matrices (left = PA runs; right = AP runs). C: Schematic of the auditory pathway from the cochlea through brainstem to cortex (https://osf.io/u2gxc/). D: Regions of interest from which functional timeseries were extracted. Top: cortical regions from FreeSurfer’s DKT atlas. Bottom: subcortical auditory regions (Sitek et al., 2019).

We found that the strongest auditory connectivity was between adjacent structures in the same hemisphere, particularly between the auditory midbrain (inferior colliculus, or IC) and thalamus (medial geniculate body, or MGB) and between core and associative auditory cortex (TTG and STG). Minimal connectivity was observed between homologous auditory structures across hemispheres. Connectivity was also present between adjacent brainstem auditory structures (cochlear nucleus [CN] and superior olivary complex [SOC]), largely bilaterally. Interestingly, despite the SOC being the primary (and earliest) decussation point in the primary auditory pathway, we only observed partial correlations between right CN and left SOC (in both data partitions), not left CN and right SOC. (See the Discussion section for potential explanations.)

### 3.2 Effect of phase encoding scheme on subcortical connectivity

Due to the anatomical location of the subcortical auditory structures—in dense, heterogeneous subcortical regions and largely adjacent to CSF—we conducted connectivity analyses separately on AP- and PA-acquired fMRI runs. We then compared the connectivity results from the two acquisition schemes. Overall connectivity patterns were highly similar between the two phase-encoding schemes, as seen in panels A and B of Figure 2. Subcortical connectivity was quite robust between the brainstem auditory regions. To quantify the similarity between the results in PA and AP acquisitions (plotted in Figure 3A), we computed the Pearson correlation coefficient *r* between the estimated partial correlations. For the proposed ARMGCGM, *r* = 0.940 with *p*- value < 0.001. To assess the similarity in sign of connectivity between acquisition schemes, we computed the Jaccard dissimilarity on the signed off-diagonals of each adjacency matrix. For the proposed ARMGCGM, the Jaccard dissimilarity was 0.231. Additionally, we computed the Euclidean distance between partial correlations in the two acquisition schemes, compared it with some selected approaches from the literature and report the results in Table 1. These results indicate strong consistency between the findings in the two different acquisition schemes.

**Table 1.** Several measures of dissimilarities between the functional connectivity networks (using full and partial correlations) estimated in each of the acquisition schemes are reported here. We provide results for all the correlation-based functional connectivity analyses, viz. ARMGCGM, full correlation, Glasso and PFA. Note that lower values of Jaccard dissimilarity and Euclidean distance implies better method.

| Method | Pearson's Correlation Coefficient | Jaccard Dissimilarity | Euclidean Distance |
| --- | --- | --- | --- |
| ARMGCGM | 0.940 ($p$-value < 0.001) | 0.231 | 0.262 |
| Full correlation | 0.811 ($p$-value < 0.001) | 0.338 | 1.016 |
| Glasso | 0.749 ($p$-value < 0.001) | 0.414 | 0.959 |
| PFM | 0.747 ($p$-value < 0.001) | 0.379 | 0.975 |

**Figure 3.**
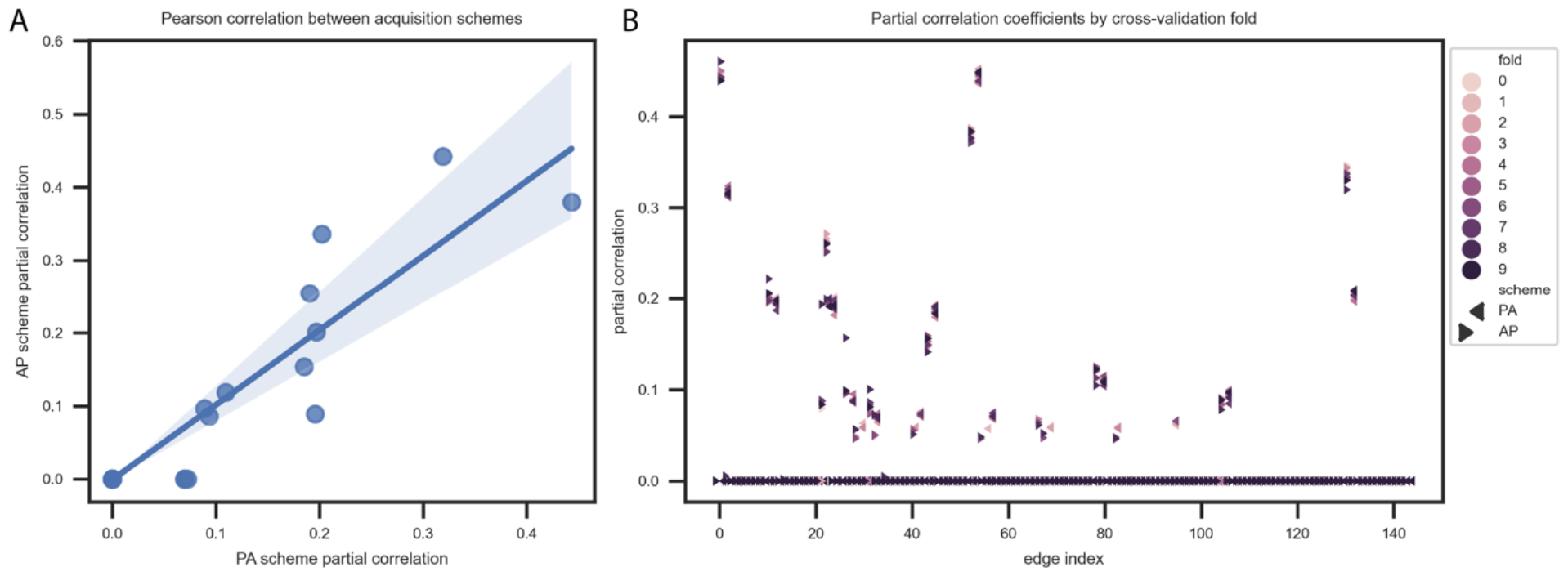
A: Pearson correlation of partial correlation values between posterior-to-anterior and anterior-to-posterior phase-encoding acquisition schemes across all edges in the graph (region-to-region connections). B: Partial correlation coefficients across cross-validation folds (10 folds for each of the two phase-encoding acquisition schemes).

### 3.3 Cross-validation of partial correlations

To assess the stability of our proposed ARMGCGM, we ran leave-10%-out cross-validation by removing 10% randomly selected subjects in each fold. We repeated this for each phase encoding scheme for a total of 20 folds; partial correlation coefficients for each fold are presented in Figure 3B. Across all folds (and both phase encoding schemes), the results are very highly similar (intraclass correlation of edgewise partial correlations across cross-validation folds = 0.991).

### 3.4 Comparison with existing approaches

We compared with three standard approaches in the literature: (1) correlation analysis between the ROIs (Biswal *et al*., 1995; Cordes *et al*., 2000; Fox and Raichle, 2007; Lowe *et al*., 1998; Smith, Vidaurre, *et al*., 2013); and partial correlation analyses using (2) the graphical lasso (Friedman *et al*., 2008) and (3) the precision factor model (Chandra *et al*., 2021). In all comparisons, we did separate analyses for each of the acquisition schemes.

#### Comparison 1:Full correlation approach

Here we study the marginal correlation between the ROIs. Letting Letting ρ_*j,j*’_ denote the correlation between ROIs *j* and *j*’ in resting state we test

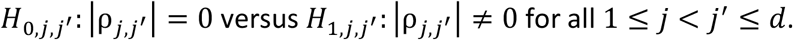

For each acquisition scheme, we first concatenate the timeseries across all runs and individuals. Then we perform *t*-tests for correlation for all (*j, j* ’) pairs followed by the Benjamini-Hochberg false discovery rate (FDR) correction (Benjamini and Hochberg, 1995) for multiplicity adjustment and control the FDR at level 0.10. Traditionally, full correlations are used to measure functional “connectivity” in resting state fMRI studies (Biswal *et al*., 1995; Cordes *et al*., 2000; Lowe *et al*., 1998). In Figure 4 we provide the correlation graphs and correlation matrices separately for each acquisition schemes. We find the correlation graphs to be much denser compared to the partial correlation graphs presented in Figure 2. Unlike in the ARMGCGM method, we observed negative values when using full correlations (particularly in the PA acquisition scheme), although the negative correlations are generally closer to 0 than the positive correlations are. In this full correlation approach, the similarity between runs split by data acquisition scheme was characterized by Pearson’s *r* of 0.811 (*p*-value < 0.001) and a Euclidean distance of 1.016. The Jaccard dissimilarity of the signed adjacency matrix was 0.338.

**Figure 4.**
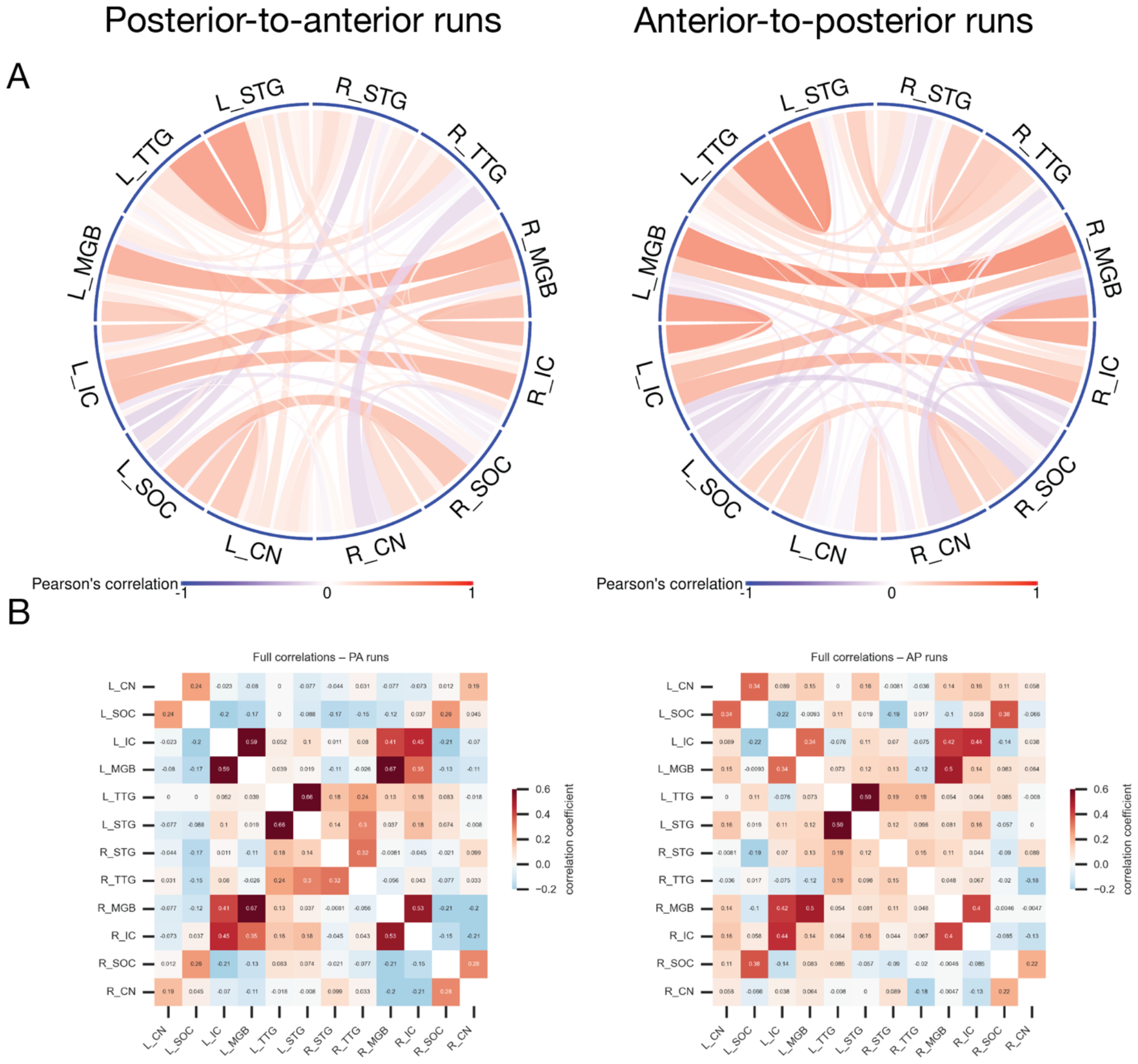
A: Full correlation connectivity in data acquired with posterior-to-anterior (PA; left) and anterior-to-posterior (AP; right) phase-encoding directions using t-tests. Positive (negative) associations are represented by red (blue) links, their opacities being proportional to the corresponding association strengths. The link widths are inversely proportional to the number of edges associated with the corresponding nodes. B: The same results as (A), viewed as adjacency matrices (left = PA runs; right = AP runs).

#### Comparison 2: Partial correlations with Glasso approach

We first consider the graphical lasso (Glasso) approach (Friedman *et al*., 2008) as another alternative choice for partial correlation based conditional graph estimation. Glasso assumes iid data from a multivariate Gaussian distribution (i.e., without any correction for autocorrelation) with 𝓁_1_(·) penalty on the precision matrix. In this analysis we concatenated the 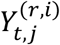 values across (*r,i*) and created a (*TR*) × *d* matrix, say ***Y***, for each acquisition scheme, and applied the Glasso model on ***Y***. We use 10-fold cross-validation to choose the optimal penalty parameter. The Glasso approach provides a point estimate of the sparse precision matrix and hence the functional connectivity network. In panels A and B of Figure 5, we plot the connectivity graphs and weighted adjacency matrices, respectively, separately for each acquisition scheme.

**Figure 5.**
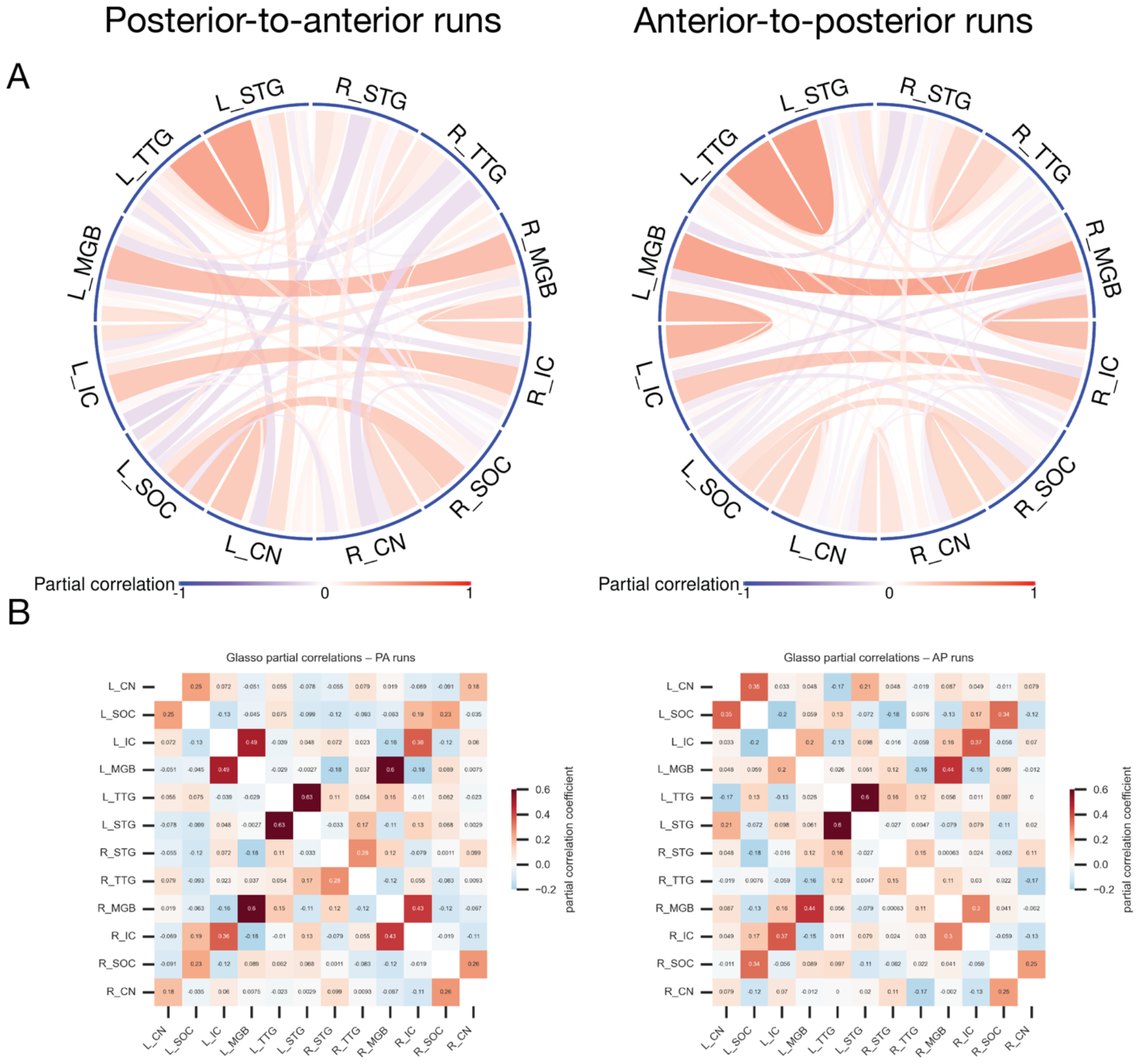
A: Partial correlation connectivity in data acquired with posterior-to-anterior (PA; left) and anterior-to-posterior (AP; right) phase-encoding directions using the Glasso approach. Positive (negative) associations are represented by red (blue) links, their opacities being proportional to the corresponding association strengths. The link widths are inversely proportional to the number of edges associated with the corresponding nodes. B: The same results as (A), viewed as adjacency matrices (left = PA runs; right = AP runs).

These functional connectivity graphs in Figure 5 are much denser compared to the estimates obtained by our proposed ARMGCGM in Figure 2. To quantify the robustness and stability of the Glasso graphs in this application, we compute Pearson correlation coefficient and Euclidean distance between the estimated partial correlations, and the Jaccard dissimilarity between the signs of the adjacency matrices in PA and AP acquisitions in the same manner as we did for ARMGCGM elaborated in Section *3*.*2*. We reported the values in Table 1. Figure 5 indicates that the strong positive correlations are consistent across the acquisition schemes. However, substantial discrepancy can be observed for the weak edges, particularly for the negative correlations. This is a common phenomenon for graphical models if the Gaussian assumption is made on non-Gaussian distributed data; see, e.g., Section 5, example 1(a) in (Guha *et al*., 2020).

#### Comparison 3: Partial correlations with PFM

In this analysis we concatenated the 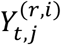 values across (*r,i*) and created a (*TR*) × *d* matrix, say ***Y***, for each acquisition scheme, and applied the PFM on ***Y***. Similar to Glasso, the PFM also assumes iid data from a multivariate Gaussian distribution and does not correct for temporal autocorrelations in the data. We infer on the graph using the Bayesian multiple comparison technique described in the Supplementary Materials. Results are provided in Figure 6. The connectivity graphs majorly differed with our ARMGCGM results, with the PFA model-derived graph being much denser and including more (weakly) negative edges. In the top panel of Figure 7 we plot the marginal Gaussian fits on the histograms of some BOLD signals. Clearly the simple Gaussian assumption does not hold here and a more sophisticated approach like ours is required.

**Figure 6.**
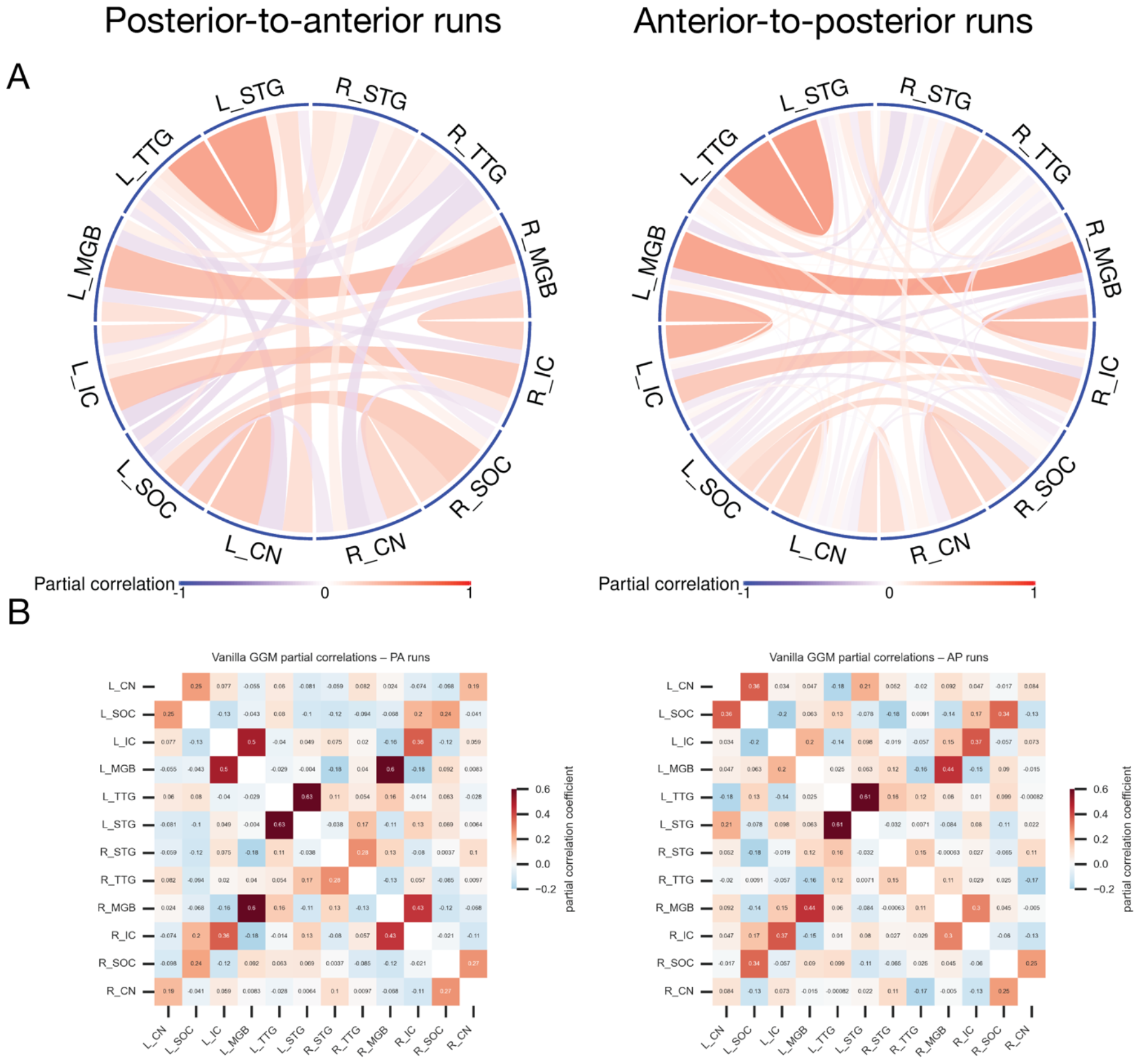
A: Partial correlation connectivity in data acquired with posterior-to-anterior (PA; left) and anterior-to-posterior (AP; right) phase-encoding directions using the PFA approach (comparison 3). Positive (negative) associations are represented by red (blue) links, their opacities being proportional to the corresponding association strengths. The link widths are inversely proportional to the number of edges associated with the corresponding nodes. B: The same results as (A), viewed as adjacency matrices (left = PA runs; right = AP runs).

**Figure 7.**
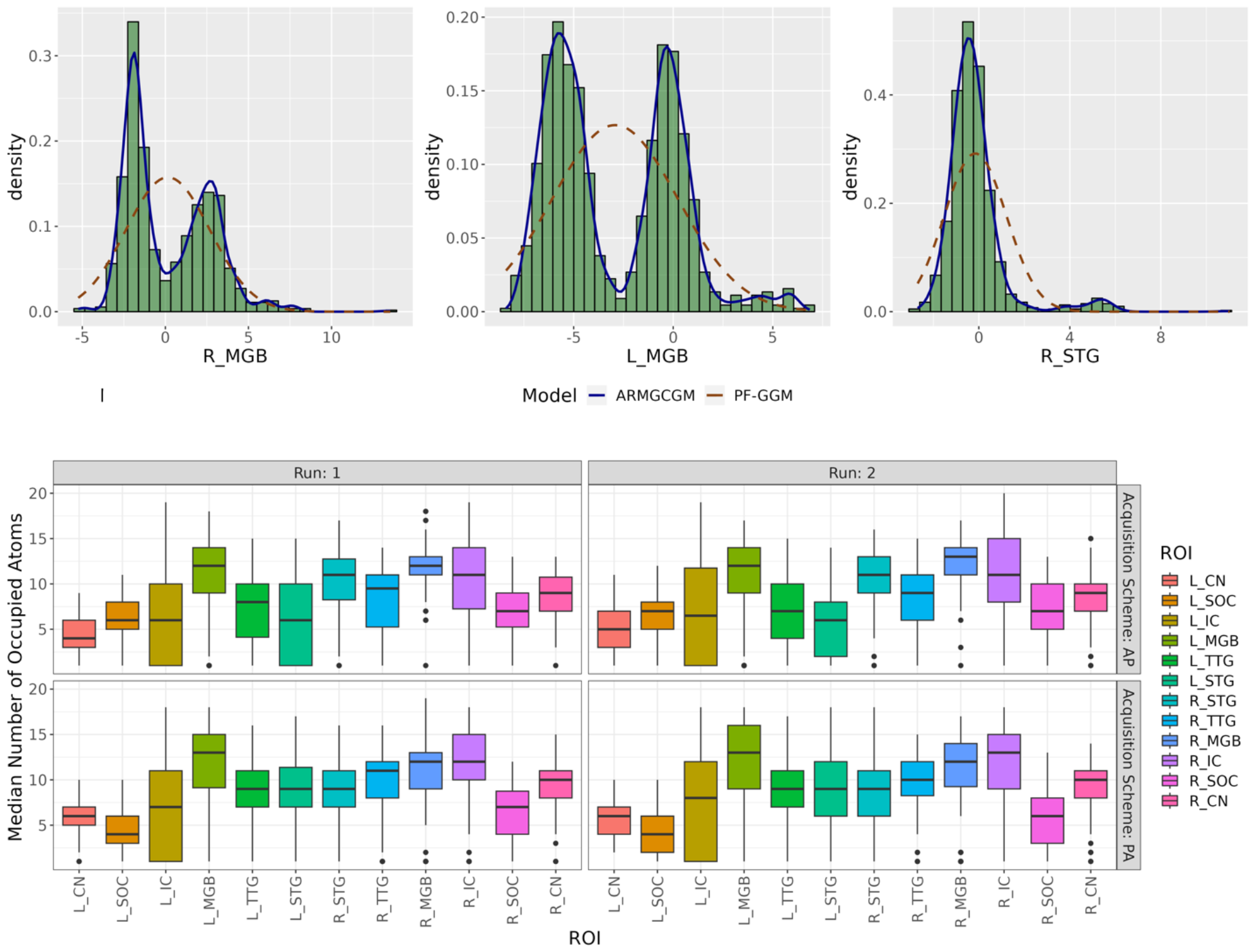
Marginal fits for the location-scale mixture of Gaussian distributions: The sample histograms in the top panel in green showed that distribution of the ROIs substantially deviates from Gaussian assumption. The blue lines were the respective fitted densities corresponding to the mixture model in equation (4). These figures indicated excellent goodness-of-fit. The dashed brown lines corresponded to marginal Gaussian fits of the vanilla PFM which were evidently underfitted. In the bottom panel we plotted the histogram of the number of occupied clusters across subjects corresponding to the mixture model indicating that our model specifications are adequate.

To assess the robustness and stability of the PFM, we computed Pearson correlation coefficient and Euclidean distance between the estimated partial correlations, and the Jaccard dissimilarity between the signs of the adjacency matrices in PA and AP acquisitions in the same manner as we did for ARMGCGM elaborated in Section *3*.*2*. We reported the values in Table 1, where we see that PFM exhibits more consistency than Glasso but ARMGCGM performed best.

### 3.5 Fit of the autoregressive matrix-Gaussian copula graphical model

#### Density fits

We studied the goodness-of-fit of the proposed ARMGCGM. In the top panel of Figure 7, we plotted the sample histograms and the corresponding fitted marginal densities for some selected ROIs. The sample histograms strongly indicate the distribution of the data to substantially deviate from Gaussian distributions, including some with multiple well-separated modes. Figure 7 also shows that our flexible location-scale mixture of Gaussians fit the data very well, even for the most complicated distributions.

Note that finite mixture models with reasonably large number of mixture components can approximate nonparametric Dirichlet process mixture models (Ishwaran and Zarepour, 2002b, 2002a). To validate whether the number of mixtures (*K* = 20) in model (4) is adequate, we computed the median number of non-empty clusters across MCMC samples for each subject and ROI in each run. In the bottom panel of Figure 7 we plotted the histograms of the medians across the subjects. As the number of non-empty clusters are smaller than ***K*** consistently across all setups yielding excellent fits for complicated distributions, we conclude that our model specifications are adequate.

#### Autocorrelation corrections

From Section 2.3 recall that 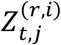’s were the transformed BOLD signal timeseries in the Gaussian space and 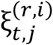’s were the autocorrelation corrected timeseries. To check for autocorrelation corrections using model (*1*), we plotted the partial autocorrelations between 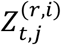’s and 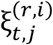’s across time for all ROIs. Since the ***Z*** and ***ξ*** values vary across MCMC iterations, we used their posterior expectations in this analysis. In Figure 8 we plot the partial autocorrelations for a randomly selected individual in run 1 of posterior-to-anterior acquisition scheme; the blue horizontal dashed lines represent the 5% band. The figure clearly indicates that the autoregressive model of order *L* = 5 corrects for the autocorrelations. As we obtained very similar results for other runs and acquisition schemes, we omit those in the paper.

**Figure 8.**
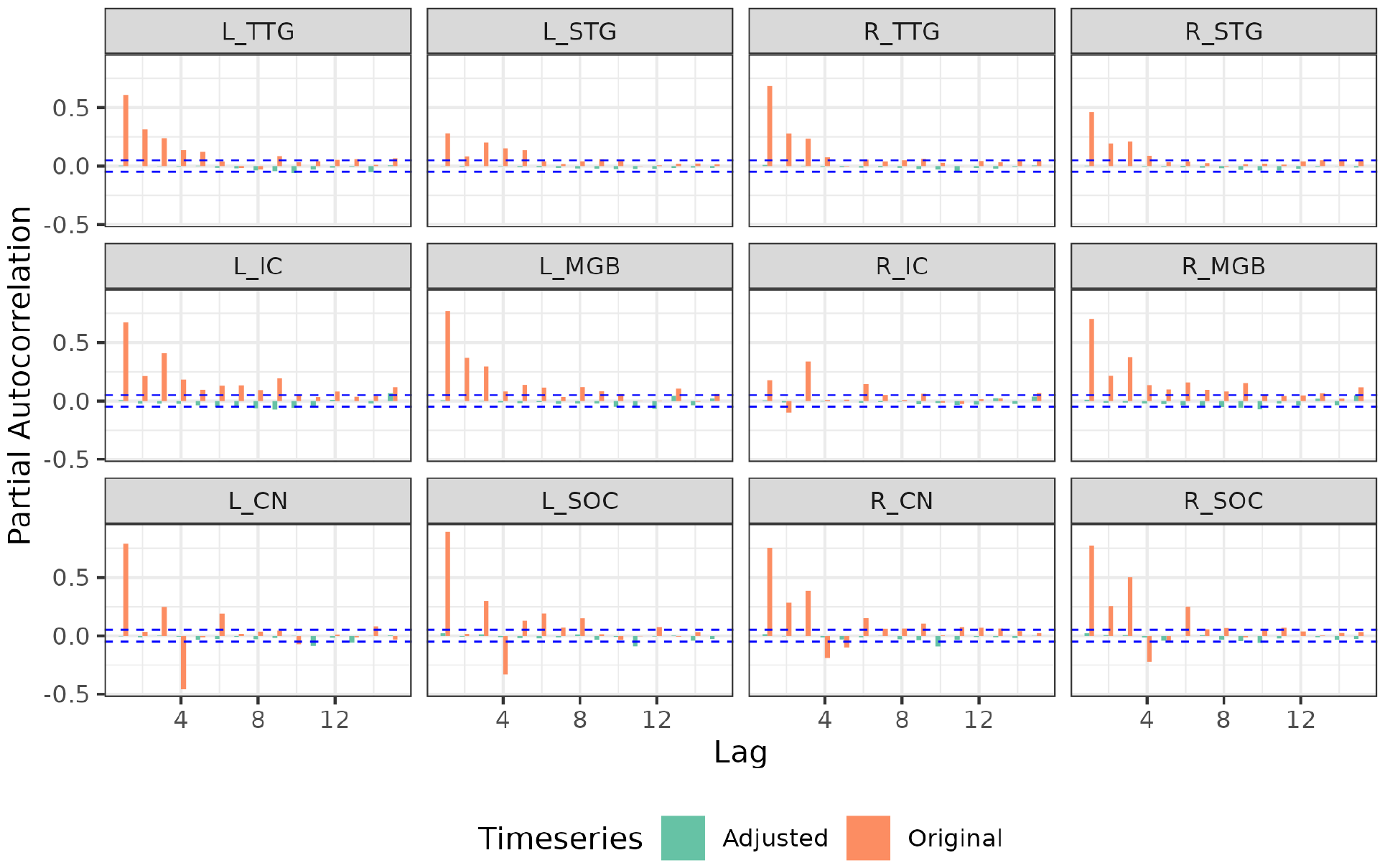
Partial autocorrelation plots of the fMRI timeseries for a randomly selected subject in the posterior-to-anterior acquisition scheme across the ROIs before (orange) and after (green) autocorrelation corrections; the horizontal dashed blue lines represent the 5% band. The plot indicates that autocorrelations are corrected in our model.

## 4. Discussion

The mammalian auditory pathway consists of a series of obligatory and interconnected subcortical and cortical brain structures. Assessing the connectivity of the human subcortical auditory structures has been limited due to methodological challenges of non-invasive imaging of the deep, small structures. Recent acquisition and analytical advances enable finer grained investigations of connectivity throughout the brain, including the brainstem. In this paper, we validated a novel an autoregressive matrix Gaussian copula graphical model to estimate functional auditory connectivity patterns from a publicly available high-resolution resting state functional MRI dataset. Using partial correlations (as opposed to full correlations) allowed us to identify specific relationships between nodes in a connectivity graph by removing shared variance across nodes (Supplementary Figures S1 and S2). We found highly consistent connectivity patterns between adjacent auditory brain regions along the auditory pathway that demonstrate the efficacy of our connectivity method as well as the potential for functional connectivity investigations of the subcortical auditory system. Below, we separately discuss our novel scientific findings and our novel contributions to the statistics literature.

### Novel contributions to the human auditory neuroscience literature

To date, there have been only limited applications of functional MRI methods to study subcortical auditory connectivity (Berlot *et al*., 2020; Hofmeier *et al*., 2018). Using our novel ARMGCGM approach in the present study, we found strong partial correlations between cochlear nucleus (CN) and superior olivary complex (SOC) bilaterally using resting state functional MRI data. Most interestingly, we observed contralateral CN–SOC connectivity between right CN and left SOC (and in both data acquisition schemes), fitting the ground truth primary auditory pathway crossing from left to right (and vice versa) between CN and SOC (Barnes *et al*., 1943; Schofield, 1994). These functional connectivity patterns between CN and SOC have not been previously observed in human auditory brainstem in vivo but follow our understanding of the mammalian primary auditory pathway based on research in animal models (Barnes *et al*., 1943; Doucet and Ryugo, 2003; Harrison and Irving, 1966). Principally, auditory information that is transduced by the cochlea of each ear is transmitted via the cochlear nerve to the cochlear nucleus, the first stage of the central auditory pathway, on each side of the brainstem. In the primary auditory pathway, the lemniscal anteroventral subdivision of the cochlear nucleus enhances the fine temporal precision of incoming auditory signals (Pickles, 2015). From there, auditory signals are passed to both the ipsilateral and contralateral SOC for further auditory processing, including spatial localization (Moore, 2000). The SOC is comprised of multiple distinct subdivisions, which receive ipsilateral and contralateral connections from cochlear nucleus to varying degrees (Pickles, 2015), aligning with our overall bilateral connectivity results between CN and SOC.

While we observed consistent right CN and left SOC connectivity, it is unclear why similar patterns were not observed between left CN and right SOC. One contributor is the lower signal-to-noise ratio in fMRI data from the lower brainstem. Paired with the small size of each of the brainstem auditory nuclei, we may still be at the edge of what is detectable using present functional connectivity methods. Additionally, this analysis was conducted on “resting state” fMRI data, during which no auditory stimuli of interest were presented or overt tasks were conducted. Resting state fMRI connectivity in the cochlear nucleus and superior olivary complex has not been examined in the previous literature to our knowledge; it is possible that sound-evoked connectivity methods would evoke greater functional connectivity, particularly in these earliest stages of the auditory pathway. Further, ipsilateral connections (i.e., between left CN and left SOC and between right CN and right SOC) may be artifactually stronger due to their close physical proximity. Even with relatively high 1.05 mm spatial resolution 7T fMRI data, CN and SOC on each side are only separated by a few voxels. These regions are thus at increased likelihood of sharing temporal fluctuations due to partial volume effects, wide point-spread functions, spatial dependence, or other as-yet-unsolved fMRI confounds that are particularly acute in the lower brainstem.

Moving up the primary auditory pathway, we observed significant partial correlations between ipsilateral inferior colliculus (IC) in midbrain and medial geniculate body (MGB) of the thalamus in both hemispheres and in both phase-encoding schemes. Inferior colliculus is a major convergence point in the auditory system, with the lemniscal IC subdivision being thought to convert distinct auditory features into discrete auditory objects for the first time in the auditory pathway. MGB continues the refinement of auditory objects via direct lemniscal connections from IC as well as rich corticofugal connections from auditory cortex to non-lemniscal MGB subdivisions (Pickles, 2015). Interestingly, we found strong partial correlations between left and right MGB in both datasets. Although not directly connected by large white matter bundles, left and right MGB are expected to process auditory information from IC at similar levels of abstraction. Thus, partial correlations may reflect indirect but shared neural mechanisms of auditory processing in the thalamus.

We did not observe IC partial correlation connectivity with either brainstem or cortical structures. IC is a key hub in the auditory system, receiving bottom-up sensory information as well as top-down modulating signals from auditory cortex and other brain regions. The lack of partial correlation connectivity with IC may be due to strong IC subdivision-specific functionality, with IC core primarily serving an ascending lemniscal role and dorsal and external IC having top-down and non-lemniscal functions. Averaging over these subdivisions may obfuscate specific connectivity patterns. Alternatively, our results may suggest that it does not have a specialized relationship with any one region beyond MGB but rather integrates and transforms auditory and other neural signals.

Finally in auditory cortex, transverse temporal gyrus (TTG)—the location of primary auditory cortex—was strongly connected with ipsilateral superior temporal gyrus (STG), which contains secondary and associative auditory cortices. Primary auditory cortex receives direct input from lemniscal MGB and is the last auditory structure with fine-grained tonotopicity (Pickles, 2015). In humans, STG is hierarchically structured, with portions further away from primary auditory cortex having increasingly wider temporal integration windows (Hamilton *et al*., 2018; Norman-Haignere *et al*., 2022) and greater categorical specificity (Bhaya-Grossman and Chang, 2022; Feng *et al*., 2021; Hamilton *et al*., 2020; Keshishian *et al*., 2023; Nourski *et al*., 2018; Pernet *et al*., 2015; Rauschecker and Tian, 2000; Rupp *et al*., 2022). While invasive recordings from human STG suggest a potential direct connection between MGB and posterior STG (Hamilton *et al*., 2021), we found mixed evidence for such a direct pathway in ourOur partial correlation data (in one hemisphere in only one of the data partitions). Our partial correlation functional connectivity results align with a vast literature demonstrating information flow between primary and non-primary auditory cortex (for review, see (Hackett, 2011). The lack of contralateral partial correlation approach measures the correlation between the concerned ROIs after filtering out the indirect effects of the remaining ROIs. Complementarily, some literature suggests that left and right auditory cortex process auditory information at distinct timescales and levels of abstraction (Güntürkün *et al*., 2020; Hickok and Poeppel, 2007; Zatorre *et al*., 2002), with left auditory cortex being uniquely tuned to rapidly changing temporal information—such as the phonetics of speech sounds—while right auditory cortex is more sensitive to slower changes in the spectral domain, particularly for speech prosody as well as music.

In comparison to our ARMGCGM partial correlation approach, we computed full correlations in the same network. Connections were much denser in the full correlation approach, aligning with the rich interconnectedness of the auditory system (Pickles, 2015). Unlike with partial correlations, which highlighted hierarchical connections between adjacent notes along the auditory pathway, we observed positive full correlations between all auditory cortical regions, regardless of hemisphere. Additionally, we found a strong positive subnetwork including IC and MGB bilaterally, whereas many of these connections (such as between left and right IC) were absent in the ARMGCGM partial correlation analysis. Since partial correlations characterize connectivity between two nodes after filtering out the effects of the other nodes, our combined results point to widespread shared information across the auditory system (per full correlation analysis) with additional shared processing between adjacent nodes of the canonical auditory pathway (per partial correlation analysis). This suggests distinct but complementary use of full and partial correlations, with full correlation analysis identifying a rich network of interconnected nodes, while partial correlations are sensitive to strong node-to-node connections.

### Novel contributions to the graphical model literature

In this article, we developed an autoregressive matrix-Gaussian copula graphical model (ARMGCGM) for non-Gaussian distributed data with temporal autocorrelation, the problem of estimating brain connectivity patterns from resting state fMRI data being the motivating problem. The ARMGCGM first uses higher order autoregressive models to capture the temporal dependence in the time series for each brain region of interest, then uses flexible location-scale mixtures of Gaussians for modeling component wise residual marginal distributions associated with different regions, and finally uses a Gaussian copula to capture the dependence across the different regions. The ARMGCGM allows borrowing of information across subjects to infer on a common connectivity graph while appropriately taking into account subject and run-specific variability via flexible mixture models. We leverage recent advances on modeling precision matrices via a flexible but computationally efficient low-rank-diagonal decomposition method that not only allows efficient exploration of the posterior space for estimating the connectivity graph but also enables easy assessment of associated uncertainty. Compared to alternative approaches to exploiting partial correlations to estimate connectivity graphs, our proposed ARMGCGM method produces results that are more consistent across the acquisition schemes with respect to multiple metrics. Additionally, the results remain highly stable in a leave-10%-out cross-validation. Considering the low signal to noise ratio in BOLD signals from deep small auditory structures, our results demonstrate the sensitivity and specificity of our model to neurobiologically plausible connections. Although the proposed ARMGCGM is a more nuanced approach and potentially capable of fitting arbitrary complicated distributions, it can be numerically expensive compared to simplistic parametric models. However, our parallelized Markov chain Monte Carlo implementation ran across all participants and runs in just over two hours, demonstrating a feasible computation time given the size of the data.

### Comparisons to connectivity literature

Our study is the first to systematically assess connectivity across the human auditory pathway using multiple connectivity measures, with previous subcortical connectivity studies limited to full correlation analysis (Berlot *et al*., 2020; Hofmeier *et al*., 2018; Leaver *et al*., 2016; Zhang *et al*., 2015). Given the anatomical and methodological constraints with subcortical fMRI, the limited fMRI connectivity literature is not too surprising. First, the deep location of the subcortical structures places them far from MRI transmit and receive coils, limiting the signal-to-noise ratio from these regions (Miletić *et al*., 2020). Because the brainstem is relatively centrally located relative to the multiple receiver coils, accelerated acquisition techniques that are based on phase differences between receiver coils are less effective (Preibisch *et al*., 2015). Second, subcortical nuclei can be quite small, requiring higher resolution imaging protocols (which unfortunately trade off SNR in order to achieve greater spatial resolution). Third, subcortical nuclei are densely organized adjacent to nuclei with heterogeneous functions, so voxels immediately next to those containing core auditory structures could contain visual, motor, or sensory nuclei, white matter, CSF, or a combination of any of these. Ultimately, each of these constraints limits the SNR from subcortical auditory nuclei.

Constraints in human subcortical auditory research have translational consequences beyond basic science. For instance, while cochlear implants have been widely successful at providing sensory information to individuals with sensorineural hearing impairments with an intact cochlear nerve (Kral *et al*., 2019; Reiss, 2020), auditory prostheses in the central auditory system have been less successful (Lim *et al*., 2009; Shetty *et al*., 2021), due in no small part to our limited understanding of the complexity of sound representation in the ascending auditory pathway.

### Limitations and future directions

As the canonical neuroanatomy of the primary auditory pathway is consistent across individuals and well-described in the literature (Sitek *et al*., 2019), we have the *a priori* expectation of a shared auditory graph across all participants. As the goal of this study was to map a network that is strongly expected based on anatomy and non-human neurophysiology, we built a joint model that includes data from all participants. This is similar to approaches used in group independent component analysis (Calhoun *et al*., 2001) and cohort-level brain mapping (Varoquaux *et al*., 2013). However, subject-specific differences in the distributions of the BOLD signals as well as autocorrelations between successive scans can induce artifactual and noisy edges in functional connectivity graphs, as seen in the Glasso and PFMs; we take care of these issues in ARMGCGM. Nevertheless, we did not investigate differences in functional connectivity between participants in the current article. Building on ARMGCGM to explore how functional connectivity varies between individuals and groups or as a function of behavior is a priority for future work.

In general, resting state fMRI connectivity measures become more reliable with longer scans (Zuo *et al*., 2019). Measurement correlations increase as time in the scanner increases, from Pearson’s *r* = 0.82 at 9 min to *r* = 0.92 at 27 min to *r* = 0.97 at 90 min (Laumann *et al*., 2015). Others described improved intraclass correlation coefficients with datasets beyond 20 minutes and up to 50 minutes (Xu *et al*., 2016). The Midnight Scan Club group (Gordon, Laumann, Gilmore, *et al*., 2017) computed a range of network connectivity metrics and found that reliability generally required at least 30 minutes of resting state data per subject. One paper (Greene *et al*., 2020) specifically investigated functional connectivity in subcortical structures and found even longer scan requirements (up to 100 minutes) for subcortical structures due to decreased signal-to-noise ratios deeper in the brain. Additionally, primary sensory networks are among the most stable within and across participants (Gratton *et al*., 2018; Hutchison *et al*., 2013). We therefore believe it is appropriate and necessary to use datasets with longer scans of resting state data in order to investigate even static subcortical auditory connectivity.

Further, while many brain networks exhibit temporal dynamics that can tell us about mental state (Fornito *et al*., 2012) or disease ((Sakoğlu *et al*., 2010); see (Hutchison *et al*., 2013) for a review), functional connectivity within primary sensory networks are among the most stable over time (Gratton *et al*., 2018), as they share bidirectional physical connections, share contributions to the same physiological tasks, and are evolutionarily conserved across species (Hutchison and Everling, 2012). In the present work, we were interested in characterizing the stationary connectivity in the primary auditory pathway that is present at rest across individuals. Adapting time-varying dynamics into this model is a promising future direction, particularly if we are interested in higher level cognitive brain networks that vary as a function of task or mental state.

## 5. Conclusions

In this article, we validated a novel autoregressive matrix Gaussian copula graphical model for partial correlation estimation while appropriately correcting for temporal autocorrelations. Using this approach, we identified functional connectivity in the human auditory system using resting state functional MRI. Whereas a complementary approach using full correlations identified a rich network of interconnected auditory regions, partial correlations highlighted direct connections between adjacent structures along auditory pathways. In particular, subcortical connectivity was highly consistent across acquisitions, demonstrating the utility and applicability of functional connectivity methods in deep brain structures. In the future, we plan to investigate whole-brain partial correlation connectivity across sensory, motor, and higher cognitive networks using the proposed models and their relationship to behavior across individuals.

## Acknowledgements

Data were provided [in part] by the Human Connectome Project, WU-Minn Consortium (Principal Investigators: David Van Essen and Kamil Ugurbil; 1U54MH091657) funded by the 16 NIH Institutes and Centers that support the NIH Blueprint for Neuroscience Research; and by the McDonnell Center for Systems Neuroscience at Washington University.

## Ethics

Data usage adhered to the Open Access Data Use Terms from the Human Connectome Project (WU-Minn HCP).

## Data and Code Availability

Data are publicly available through the Human Connectome Project (https://www.humanconnectome.org/study/hcp-young-adult). Analysis code is available in the Supplementary Materials.

## Author Contributions

Noirrit Chandra: conceptualization (equal); formal analysis (lead); software (lead); visualization (equal); writing – original draft (equal); writing – review and editing (equal).

Kevin R. Sitek: conceptualization (equal); formal analysis (equal); visualization (equal); writing – original draft (lead); writing – review and editing (equal).

Bharath Chandrasekaran: conceptualization (equal); writing – review and editing (equal). Abhra Sarkar: conceptualization (equal); writing – review and editing (equal).

