## Supplemental Material for "Functional connectivity across the human subcortical auditory system using an autoregressive matrix-Gaussian copula graphical model approach with partial correlations"

### S1. Supplementary Materials

#### S1.1 Full vs Partial Correlation

Here, we illustrate the difference between full and partial correlation analyses in a stylized example. We consider the following full correlation matrix in the left panel of Supplementary Figure 1 and on the right panel on the same figure, we show the marginal dependency graph. We can see that marginally all of the nodes are dependent on each other.

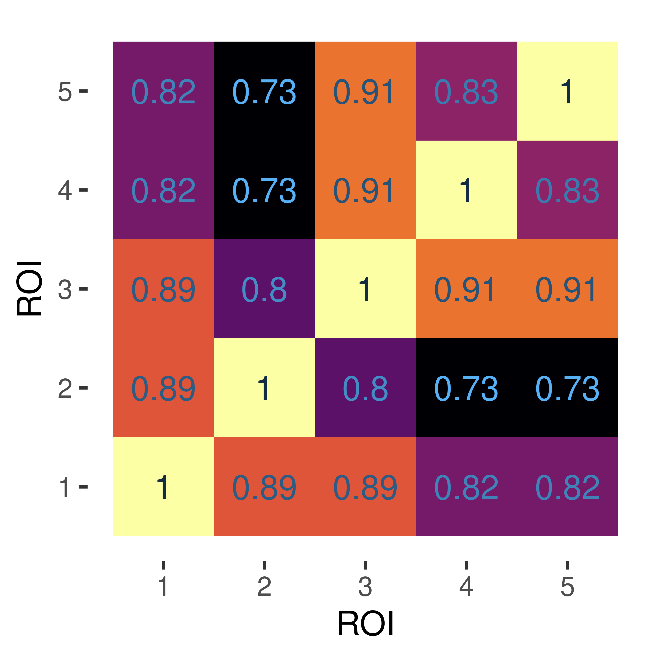

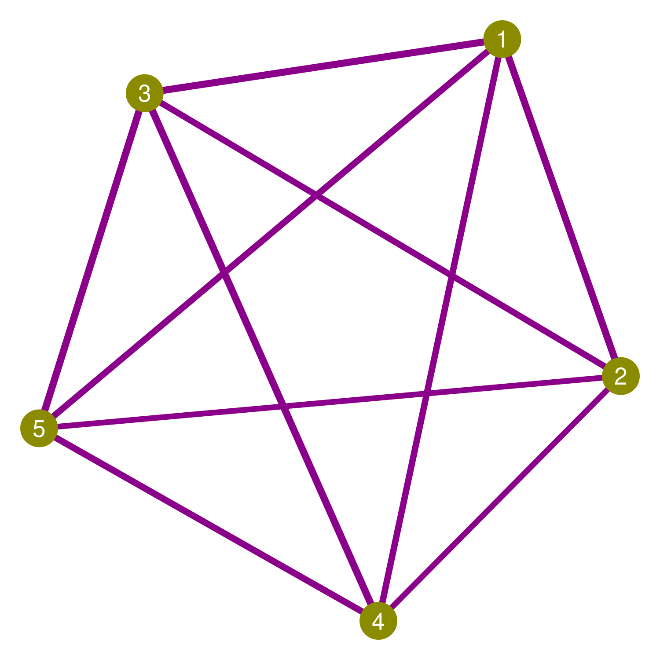

Figure 1 Simple correlation matrix (left) and marginal dependency graph (right).

We calculate the partial correlation matrix and show in the left panel of Supplementary Figure 2. Partial correlations represent the dependence between two variables after removing the effects of the other variables. For example, 0.58 is the partial correlation between region of interest (ROI)-5 and ROI-3 after removing the effects of ROIs 1, 2 and 4. On the left panel of Figure 2 we show the corresponding conditional dependency graph. Even though the marginal graph is dense, we see that the conditional dependence graph is very sparse.

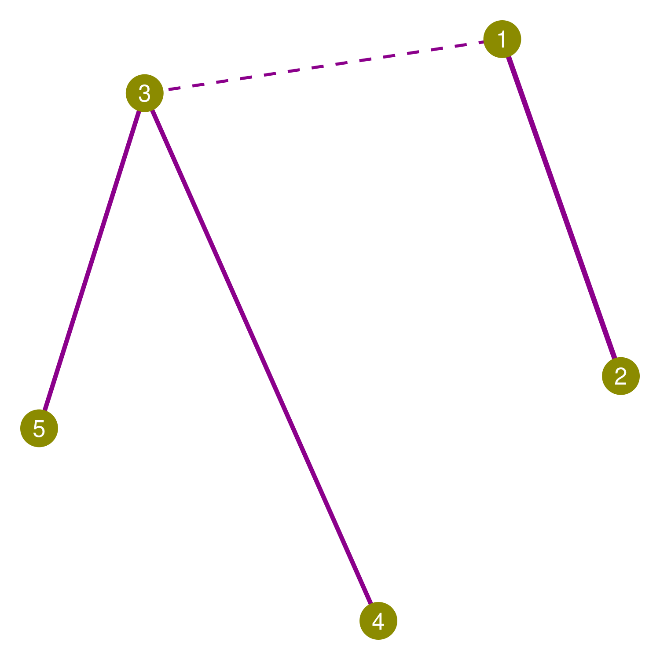

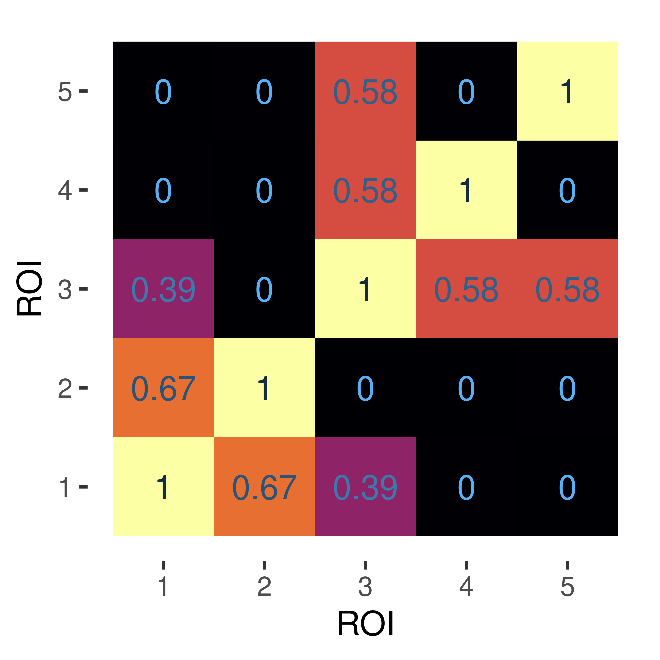

Figure 2 Partial correlation matrix (left) and conditional dependency graph (right).

#### S1.2 Posterior Computation

For each ROI, we have timeseries of length $T$ from each of the $n$ subjects. Our Gibbs sampler algorithm to draw samples from the posterior distribution cycles through the following steps.

Step 1 **Update the mixture model parameters:** Note that the mixture model in equation (4) of the main paper can be equivalently represented as

$$Y_{t,j}^{\left( r,i \right)}\mid c_{t,j}^{\left( r,i \right)}=h\sim\text{N}\left( \mu_{h,j}^{\left( r,i \right)},\sigma_{h,j}^{2\left( r,i \right)} \right),\quad\Pr\left( c_{t,j}^{\left( r,i \right)}=h \right)=\pi_{h,j}^{\left( r,i \right)}\text{ for all }t=1,\ldots,T,$$

(1)

where $c_{t,j}^{\left( r,i \right)}$s are latent cluster membership indicators. We update these latent variables and the parameters in the following steps.

**Parallel for** $i=1,\ldots,N$

for $j=1,\ldots,d$ and $r=1,\ldots,R$

i) for $t=1,\ldots,T$ and $h=1,\ldots,K$, update $c_{t,j}^{\left( r,i \right)}$ by sampling $\Pr\left( c_{t,j}^{\left( r,i \right)}=h \right)\propto\pi_{h,j}^{\left( r,i \right)}\text{N}\left( \mu_{h,j}^{\left( r,i \right)},\sigma_{h,j}^{2\left( r,i \right)} \right)$;
ii) for $h=1,\ldots,K$ define the set $S_{h,j}^{\left( r,i \right)}=\left\{ t:c_{t,j}^{\left( r,i \right)}=h \right\}$ and the quantities $N_{h,j}^{\left( r,i \right)}=$ cardinality of $S_{h,j}^{\left( r,i \right)}$, $\hat{\nu}=\nu_{0}+N_{h,j}^{\left( r,i \right)}$, $\hat{a}=a_{0}+\frac{N_{h,j}^{\left( r,i \right)}}{2}$, $\overline{Y}_{h,j}^{\left( r,i \right)}=\frac{1}{N_{h,j}^{\left( r,i \right)}}\sum_{t\in S_{h,j}^{\left( r,i \right)}} Y_{t,j}^{\left( r,i \right)}$, $\hat{\mu}=\frac{\nu_{0}\mu_{0}+N_{h,j}^{\left( r,i \right)}\overline{Y}_{h,j}^{\left( r,i \right)}}{\hat{\nu}}$, $\hat{b}=b_{0}+\frac{1}{2}\sum_{t\in S_{h,j}^{\left( r,i \right)}} \left( Y_{t,j}^{\left( r,i \right)}-\overline{Y}_{h,j}^{\left( r,i \right)} \right)^{2}+\frac{N_{h,j}^{\left( r,i \right)}\nu_{0}}{2\hat{\nu}}\left( \overline{Y}_{h,j}^{\left( r,i \right)}-\mu_{0} \right)^{2}$. Then sample $\left( \mu_{h,j}^{\left( r,i \right)},\sigma_{h,j}^{2\left( r,i \right)} \right)\sim\text{NIG}\left( \hat{\mu},\hat{\nu},\hat{a},\hat{b} \right)$;
iii) sample $\left( \pi_{1,j}^{\left( r,i \right)},\ldots,\pi_{K,j}^{\left( r,i \right)} \right)\sim\text{Dir}\left( N_{1,j}^{\left( r,i \right)}+\frac{\alpha_{\pi}}{K},\ldots,N_{K,j}^{\left( r,i \right)}+\frac{\alpha_{\pi}}{K} \right)$.

Step 2 **Update the autoregressive process parameters:** This cycles through the following steps
       **Parallel for** $i=1,\ldots,N$
       for $j=1,\ldots,d$ and $r=1,\ldots,R$
       (i) Calculate $Z_{t,j}^{\left( r,i \right)}=\Phi^{-1}\left\{ F_{j}^{\left( r,i \right)}\left( Y_{t,j}^{\left( r,i \right)} \right) \right\}$ where $\Phi\left( \cdot\right)$ is the CDF of a standard Gaussian distribution and $F_{j}^{\left( r,i \right)}\left( \cdot\right)$ is the CDF induced by the mixture model (*1*).
       (ii) Letting $\boldsymbol{Z}^{\left( r,i \right)}=\left( \left( Z_{t,j}^{\left( r,i \right)} \right) \right)_{T\times d}$ denote the matrix of fMRI signals corresponding to the $i$-th individual in the $r$-th run in the transformed Gaussian space, obtain ${\tilde{\boldsymbol{Z}}}^{\left( r,i \right)}=\boldsymbol{Z}^{\left( r,i \right)}\boldsymbol{R}_{\Omega}^{\frac{\boldsymbol{1}}{\boldsymbol{2}}}$.
       (iii) Letting $\boldsymbol{z}_{j}$ denote the $j$-th column of ${\tilde{\boldsymbol{Z}}}^{\left( r,i \right)}$, we define the $T-1$ component vector $\boldsymbol{\eta}=\left( z_{2},\ldots,z_{T} \right)^{T}$ and the $\left( T-1 \right)\times L$ matrix $\boldsymbol{X}=\left( \left( z_{i-j+1} \right) \right)_{\left( T-1 \right)\times L}$with the convention $z_{t^{'}}=0$ for any $t^{'}\leq0.$ Then sample

$$\boldsymbol{\beta}_{j}^{\left( r,i \right)}=\left( \beta_{1,j}^{\left( r,i \right)},\ldots,\beta_{L,j}^{\left( r,i \right)} \right)^{T}\sim t_{2\tilde{a}_{\varsigma}}\left( \tilde{\boldsymbol{\beta}},\frac{\tilde{b}_{\varsigma}}{\tilde{a}_{\varsigma}}\boldsymbol{\Sigma}_{\beta}^{-1} \right),\quad\quad\varsigma_{j}^{-2\left( r,i \right)}\sim\text{Ga}\left( \tilde{a}_{\varsigma},\tilde{b}_{\varsigma} \right),$$

where $\tilde{a}_{\varsigma}=a_{\varsigma}+\frac{T-1}{2}$, $\boldsymbol{\Sigma}_{\beta}=\boldsymbol{X}^{T}\boldsymbol{X}+\boldsymbol{I}_{L}$, $\tilde{\boldsymbol{\beta}}=\boldsymbol{\Sigma}_{\beta}^{-1}\boldsymbol{X}^{T}\boldsymbol{\eta}$, $\tilde{b}_{\varsigma}=b_{\varsigma}+\frac{1}{2}\left( \boldsymbol{\eta}^{T}\boldsymbol{\eta}-{\tilde{\boldsymbol{\beta}}}^{T}\boldsymbol{\Sigma}_{\beta}^{\boldsymbol{-1}}\tilde{\boldsymbol{\beta}} \right)$ and a $p$- dimensional $t_{\nu}\left( \boldsymbol{x};\boldsymbol{\mu},\boldsymbol{\Sigma} \right)$ denotes the central-$t$ distribution with location vector $\boldsymbol{\mu}$ and scale matrix $\boldsymbol{\Sigma}$ and the following pdf $\frac{\Gamma\left( \frac{\nu+p}{2} \right)}{\Gamma\left( \frac{\nu}{2} \right)\left( \nu\pi\right)^{\frac{p}{2}}\left| \boldsymbol{\Sigma} \right|^{\frac{1}{2}}}\left\{ 1+{{\frac{1}{\nu}\left( \boldsymbol{x}-\boldsymbol{\mu} \right)}^{T}\boldsymbol{\Sigma}}^{-1}\left( \boldsymbol{x}-\boldsymbol{\mu} \right) \right\}^{-\frac{\nu+p}{2}}$.
Step 3 **Sample the latent variables for updating** $\boldsymbol{\Omega}$**:**
       (i) Calculate the autocorrelation corrected Gaussian values $\hat{Z}_{t,j}^{\left( r,i \right)}=\frac{1}{\varsigma_{j}^{\left( r,i \right)}}\left( \tilde{Z}_{t,j}^{\left( r,i \right)}-\sum_{t^{'}=1}^{L} \beta_{t^{'},j}^{\left( r,i \right)}\tilde{Z}_{t-t^{'},j}^{\left( r,i \right)} \right)$ and subsequently obtain $\boldsymbol{W}^{\left( r,i \right)}={\hat{\boldsymbol{Z}}}^{\left( r,i \right)}\boldsymbol{R}_{\Omega}^{\frac{1}{2}}$ where ${\hat{\boldsymbol{Z}}}^{\left( r,i \right)}=\left( \left( \hat{Z}_{t,j}^{\left( r,i \right)} \right) \right)$.
       (ii) Construct the $nTR\times d$ matrix $\boldsymbol{W}$ by concatenating the $\boldsymbol{W}^{\left( r,i \right)}$ matrices across all $r=1,\ldots,R$ and $i=1,\ldots,n$.
       (iii) For notational convenience, we let $N=nTR$ and define $\boldsymbol{W}_{i}=\left( W_{i,1},\ldots,W_{i,d} \right)^{T}$. Generate $q$-dimensional vectors $\boldsymbol{u}_{1},\ldots,\boldsymbol{u}_{N}\overset{iid}{\sim}\text{N}_{q}\left( \boldsymbol{0},\boldsymbol{P} \right)$ with $\boldsymbol{P}=\left( \boldsymbol{I}_{q}+\boldsymbol{\Lambda}^{T}\boldsymbol{\Delta}^{\boldsymbol{-1}}\boldsymbol{\Lambda} \right)$ independently from $\boldsymbol{W}_{1:N}$ and let $\boldsymbol{\vartheta}_{i}=\boldsymbol{W}_{i}+\boldsymbol{\Delta}^{\boldsymbol{-1}}\boldsymbol{\Lambda}\boldsymbol{P}^{\boldsymbol{-1}}\boldsymbol{u}_{i}$.
Step 4 **Update** $\boldsymbol{\Lambda}$**:** We have $\boldsymbol{u}_{i}=\sum_{r=1}^{d} \boldsymbol{\lambda}_{r}\vartheta_{r,i}+\boldsymbol{\varepsilon}_{i}$, where $\boldsymbol{\lambda}_{r}=\left( \lambda_{r,1},\ldots,\lambda_{r,q} \right)$ is the $r$-th row of $\boldsymbol{\Lambda}$ and $\boldsymbol{\vartheta}_{i}=\left( \vartheta_{1,i},\ldots,\vartheta_{d,i} \right)^{T}$. Define $\boldsymbol{u}_{i}^{\left( j \right)}=\boldsymbol{u}_{i}-\sum_{r\neq j} \boldsymbol{\lambda}_{r}\vartheta_{r,i}.$ Then $\boldsymbol{u}_{i}^{\left( j \right)}=\boldsymbol{\lambda}_{j}\vartheta_{j,i}+\boldsymbol{\varepsilon}_{i}.$ Conditioned on $\boldsymbol{u}_{i}^{\left( j \right)}$, $\boldsymbol{\vartheta}_{i}$ and the associated hyper-parameters, $\boldsymbol{\lambda}_{j}$'s can be updated sequentially for $j=1,\ldots,d$ from the distribution

$$\boldsymbol{\lambda}_{j}\sim\text{N}_{\text{q}}\left\{ \left( \boldsymbol{D}_{j}^{-1}+\left\| \boldsymbol{\vartheta}^{\left( j \right)} \right\|^{2}\boldsymbol{I}_{q} \right)^{-1}\boldsymbol{m}_{j},\left( \boldsymbol{D}_{j}^{-1}+\left\| \boldsymbol{\vartheta}^{\left( j \right)} \right\|^{2}\boldsymbol{I}_{q} \right)^{-1} \right\},$$

where $\boldsymbol{D}_{j}=\tau^{2}\text{diag}\left( \psi_{j,1}\phi_{j,1}^{2},\ldots,\psi_{j,q}\phi_{j,q}^{2} \right)$, $\boldsymbol{\vartheta}^{\left( j \right)}=\left( \vartheta_{j,1},\ldots,\vartheta_{j,N} \right)^{T}$ and $\boldsymbol{m}_{j}=\sum_{i=1}^{N} \vartheta_{j,i}\boldsymbol{u}_{i}^{\left( j \right)}.$

Step 5 **Update** $\boldsymbol{\Delta}$ **and** $\alpha$**:** Sample the $\delta_{j}^{2}$'s through the following steps.
       (i) Let $d_{r,-j}=\sum_{l\neq j} 1\left( \varsigma_{l}=r \right)$ and $\boldsymbol{\vartheta}^{\left( -j \right)}$ to be the collection of all $\boldsymbol{\vartheta}^{\left( l \right)}$'s, $l=1,\ldots,d$, excluding $\boldsymbol{\vartheta}^{\left( j \right)}$ and $1\left( \cdot\right)$ be the indicator function. For $j=1,\ldots,d$, sample the cluster indicators sequentially from the distribution

$$\Pr\left( \varsigma_{j}=r \right)\propto\left\{ \begin{aligned} d_{r,-j}\int\text{N}\left( \boldsymbol{\vartheta}^{\left( j \right)};0,\delta_{r}^{-2} \right)\text{d}G_{0}\left( \delta_{r}^{2}|\boldsymbol{\vartheta}^{\left( -j \right)} \right)\text{ for }r\in\text{\{}\varsigma_{l}{\text{\}}}_{l\neq j}; \\ \alpha\int\text{N}\left( \boldsymbol{\vartheta}^{\left( j \right)};0,\delta_{r}^{-2} \right)\text{d}G_{0}\left( \delta_{r}^{2} \right)\text{ for }r\neq\varsigma_{l}\text{ for all }l\neq j. \end{aligned} \right.$$

The above integrals are analytically available and involves the density of a multivariate central Student's $t$-distribution for $G_{0}=\text{Ga}\left( a_{\delta},b_{\delta} \right).$

(ii) Let the unique values in $\varsigma_{1:d}$ be $\text{\{}1,\cdots,k\text{\}}.$ For $r=1,\ldots,k$, set $d_{r}=\sum_{j} 1\left( \varsigma_{j}=r \right)$ and $V_{r}=\sum_{j:\varsigma_{j}=r} \left\| \vartheta^{\left( j \right)} \right\|^{2},$ and independently sample $\delta_{r}^{2}\sim\text{Ga}\left( a_{\delta}+Nd_{r}\text{/}2,b_{\delta}+V_{r}\text{/}2 \right)$.
(iii) Set $\delta_{j}^{2}=\delta_{\varsigma_{j}}^{2}$.
(iv) Following (West 1992), first generate $\varphi\sim\text{Beta}\left( \alpha+1,d \right)$, evaluate $\pi\text{/}\left( 1-\pi\right)=\left( a_{\alpha}+k-1 \right)\text{/}\left\{ d\left( b_{\alpha}-log\varphi\right) \right\}$ and then generate

$$\alpha|\varphi,k\sim\left\{ \begin{aligned} \text{Ga}\left( \alpha+k,b_{\alpha}-log\varphi\right)\text{ with probability }\pi; \\ \text{Ga}\left( \alpha+k-1,b_{\alpha}-log\varphi\right)\text{ with probability}1-\pi. \end{aligned} \right.$$

Step 6 **Update Dirichlet-Laplace hyperparameters:** Sample the hyper-parameters in the priors on $\boldsymbol{\Lambda}$ through the following steps.
       (i) For $j=1,\ldots,d$ and $h=1,\ldots q$ sample $\tilde{\psi}_{j,h}$ independently from an inverse-Gaussian distribution $\text{iG}\left( \tau\phi_{j,h}\text{/}\left| \lambda_{j,h} \right|,1 \right)$ and set $\psi_{j,h}=1\text{/}\tilde{\psi}_{j,h}.$
       (ii) Sample the full conditional posterior distribution of $\tau$ from a generalized inverse Gaussian $\text{giG}\left\{ dq\left( 1-a \right),2b,2\sum_{j,h} \left| \lambda_{j,h} \right|\text{/}\phi_{j,h} \right\}$ distribution.
       (iii) Draw $T_{j,h}$ independently with $T_{j,h}\sim\text{giG}\left( a-1,1,2\left| \lambda_{j,h} \right| \right)$ and set $\phi_{j,h}=T_{j,h}\text{/}T$ with $T=\sum_{j,h} T_{j,h}$.

#### S1.3 Choice of Hyperparameters

*Autoregressive model hyperparameters:* We let $\nu_{\beta}=1$ and set $\left( a_{\varsigma},b_{\varsigma} \right)$ such that *a priori* $E\left( \varsigma_{j}^{2\left( r,i \right)} \right)=1$ and V$\mathrm{ar}\left( \varsigma_{j}^{2\left( r,i \right)} \right)=5$ for all $r,i,j$ to ensure weakly informative priors. We set the order of the autoregressive models $L=5$.

*Mixture model hyperparameters:* Regarding the Dirichlet prior’s concentration parameter, we let $\alpha_{\pi}=1$. Regarding the normal-inverse gamma prior on the location-scale parameters of the mixture components, we let $\mu_{0}=0$, $\nu_{0}=0.01$, $a_{0}=250$ and $b_{0}=20$. Such hyperparameter choices imply $E\left( \mu_{h,j}^{\left( r,i \right)} \right)=0$ and $E\left( \sigma_{h,j}^{2\left( r,i \right)} \right)\approx0$ for all $r,i,j,h$. Based on previous experience on Gaussian mixture models, we observed that such choices favor a larger number of occupied clusters in the mixture model a posteriori, allowing for a more flexible fit. We set number of mixture components $K=20$.

#### Precision factor analysis model hyperparameters: In this paper we set the column-dimension $(q)$ of $\boldsymbol{\Lambda}$ $q=d$ where $d$ is the row-dimension of $\boldsymbol{\Lambda}$, i.e., the number of ROIs of interest. Since the number of ROIs is in the order of tens in this paper, we take $q=d$ to have a full-rank model.

#### Following the suggestions of (Bhattacharya et al. 2015; Chandra et al. 2021) we let $a=0.5$ and $b=2$ as the hyperparameters of the two-parameter Dirichlet-Laplace prior on $\boldsymbol{\Lambda}$. For the prior on the residual variances and the Dirichlet process concentration parameter, we set ${a_{\delta}=b_{\delta}=a}_{\alpha}=b_{\alpha}=0.10$ implying weakly informative priors.

#### S1.4 Graph Selection

Typical of Bayesian continuous shrinkage priors, exact zero estimates are not obtained even for the insignificant off-diagonal elements of $\boldsymbol{\Omega}$ for finite sample sizes. This is an artifact of continuous shrinkage priors, since the probabilities of exact zeroes are *almost surely* zero although the posterior probabilities of arbitrary sets around zeroes are very high.

We address the issue of non-zero edge selection through a multiple hypothesis testing based approach. For $i=1,\ldots,d, j=i+1,\ldots,d$ and some $\epsilon>0$, we consider testing

$$H_{0,i,j}:\left| \rho_{i,j} \right|\leq\epsilon\mathrm{versus} H_{1,i,j}:\left| \rho_{i,j} \right|>\epsilon,$$

where $\rho_{i,j}$ is the $\left( i,j \right)$-th element of ${diag\left( \boldsymbol{\Omega} \right)}^{-\frac{1}{2}} {\boldsymbol{\Omega} diag\left( \boldsymbol{\Omega} \right)}^{-\frac{1}{2}}$,the partial correlation matrix derived from $\boldsymbol{\Omega}$. Here we follow (Berger 1985) (Chapter 4, pp. 148) in replacing the point nulls $H_{0,i,j}:\rho_{i,j}=0$ by reasonable interval nulls $H_{0,i,j}:\left| \rho_{i,j} \right|\leq\epsilon$. If $H_{0,i,j}$ is rejected in favor of $H_{1,i,j}$, we conclude that there is an edge between nodes $i$ and $j$. We utilize posterior uncertainty to resolve these testing problems. Specifically, we define $d_{i,j}=1\left\{ \Pi\left( H_{1,i,j} | Y \right)>\beta\right\}$ as the decision rule which controls the posterior FDR defined as

$$FDR_{Y}=\frac{\sum_{i,j} d_{i,j}\Pi\left( H_{0,i,j} | Y \right)}{\max\left( \sum_{i,j} d_{i,j},1 \right)},$$

at the level $1-\beta$. Importantly, the decision rule also incurs the lowest false non-discovery rate (Müller et al. 2004). For a fixed $\beta$, the $FDR_{Y}$ depends on the choice of $\epsilon$. To obtain the optimal $\epsilon$, we compute the $FDR_{Y}$'s on a grid of $\epsilon$ values in $\left( 0,1 \right)$ and then set $\epsilon={\inf_{\epsilon^{'}} FDR}_{Y}\left( \epsilon^{'} \right)\leq1-\beta$. In all applications in this paper, we control $FDR_{Y}$ at the $0.10$ level of significance.

#### S1.4 Control ROI graph

To assess whether our ARMGCGM approach is sensitive to connectivity differences across brain networks, we conducted an analysis that included four control regions (two per hemisphere): pericalcarine cortex and superior frontal gyrus. Pericalcarine cortex is the anatomical location of primary visual cortex, while superior frontal gyrus was selected due to its lack of association with auditory processing in Neurosynth (https://neurosynth.org/analyses/terms/auditory/). Each of these control regions is associated with cortical networks that are distinct from networks involved in auditory processing (Yeo et al. 2011).

Because partial correlations are influenced by the nodes that are included in the graph, specific edges may change depending on the makeup of the graph. However, we found that connectivity between auditory regions did not differ notably in this graph that includes control regions. Additionally, the strongest partial correlations of each control region was with its contralateral homolog. These results were consistent between data splits (by acquisition scheme). These results suggest that partial correlations using our ARMGCGM are specific to their individual subnetworks.

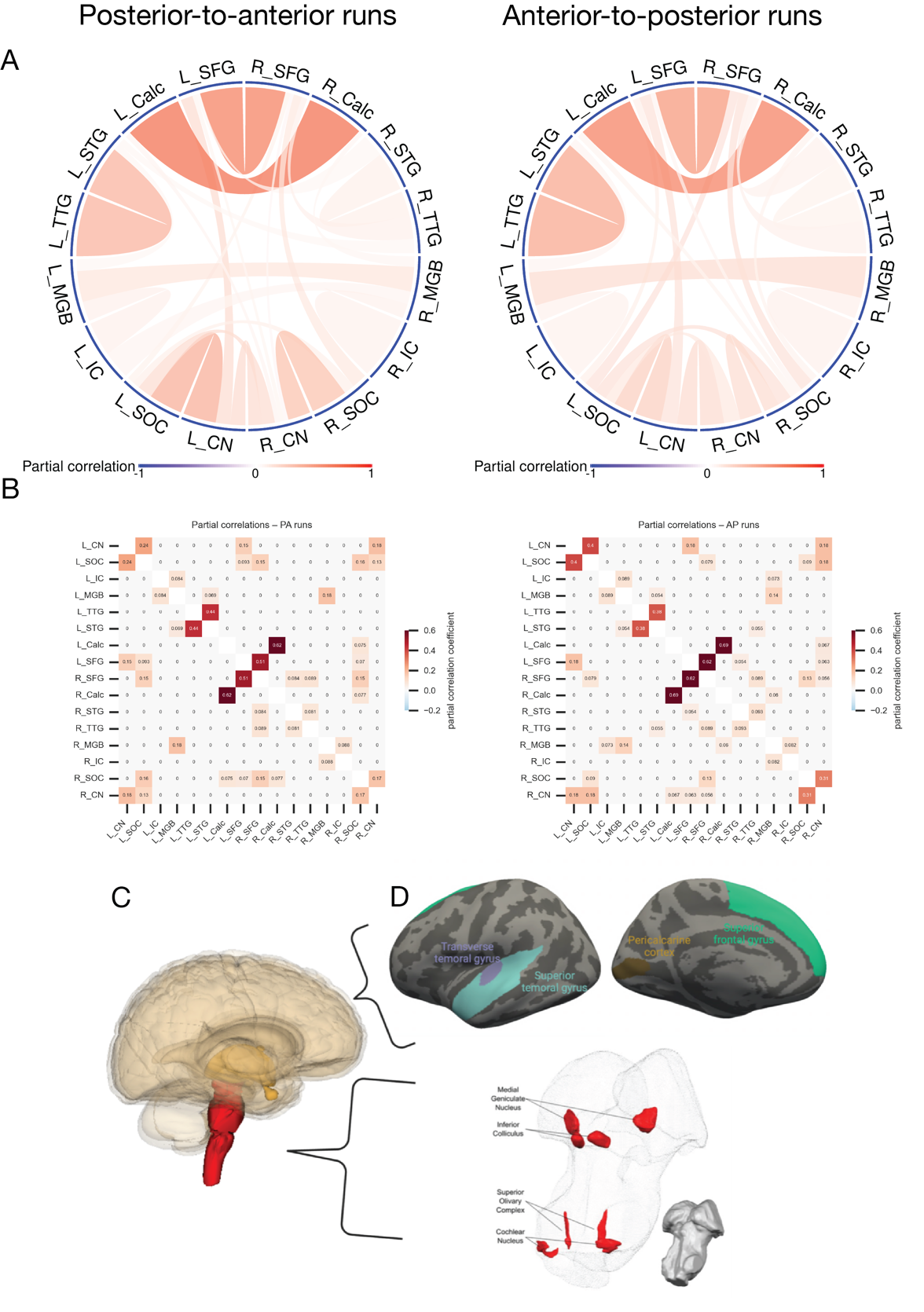

Figure 3. A: Partial correlation connectivity in data acquired with posterior-to-anterior (PA; left) and anterior-to-posterior (AP; right) phase-encoding directions using the ARMGCGM approach. Positive (negative) associations are represented by red (blue) links, their opacities being proportional to the corresponding association strengths. The link widths are inversely proportional to the number of edges associated with the corresponding nodes. B: The same results as (A), viewed as adjacency matrices (left = PA runs; right = AP runs). C: 3-dimensional view of the human brain with the brainstem highlighted in red. C: regions of interest from which functional timeseries were extracted. Top: cortical regions from FreeSurfer’s DKT atlas. Bottom: subcortical auditory regions (Sitek et al. 2019).

#### S1.5 Data and analysis workflow

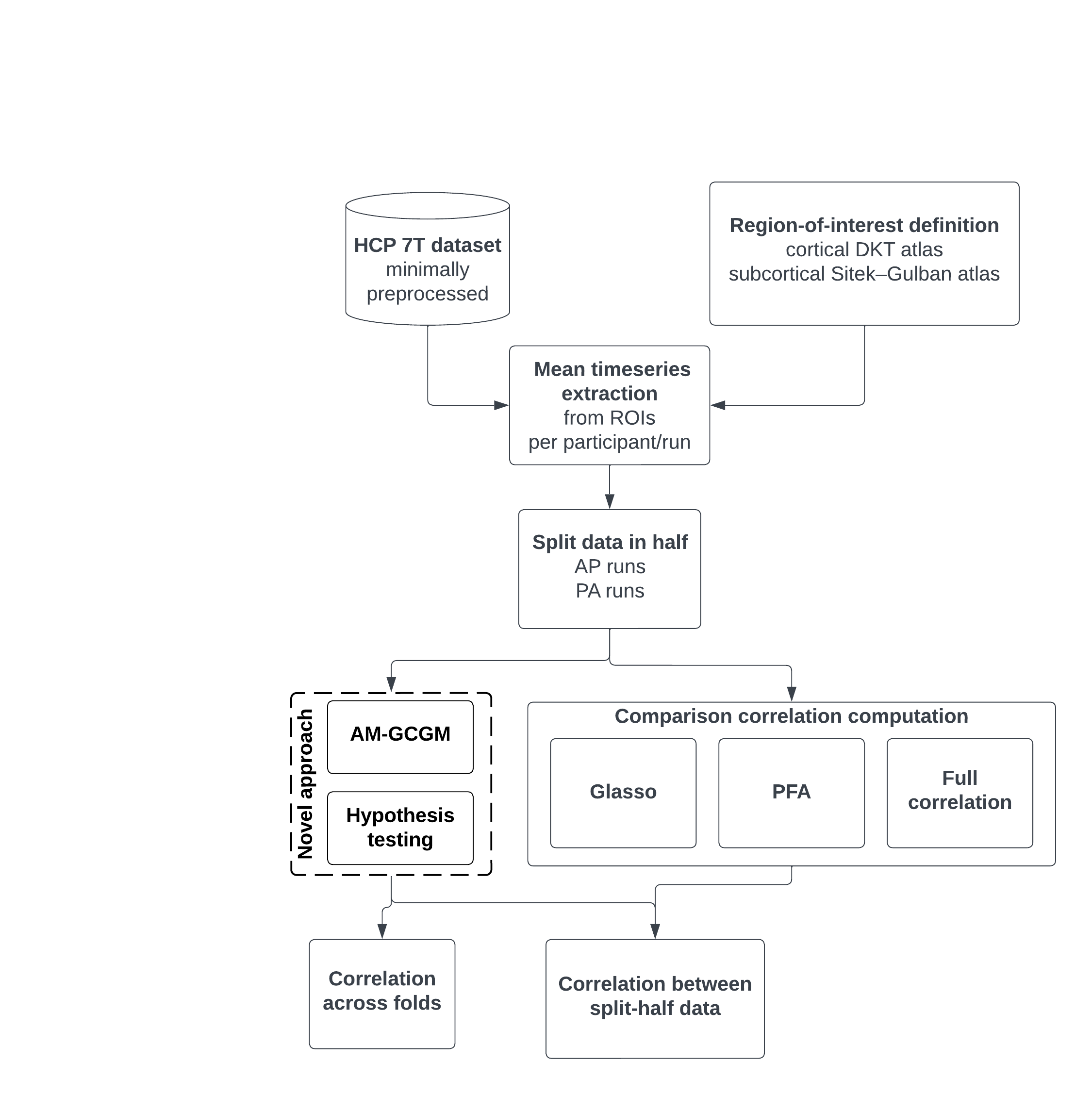

Figure 4. High-level diagram of materials and methods implemented in this manuscript.
