## Supplementary material for "Functional connectivity across the human subcortical auditory system using an autoregressive matrix-Gaussian copula graphical model approach with partial correlations": Code: rs_atlas.py

```
import os
from glob import glob

from nipype import config
config.set('execution', 'remove_unnecessary_outputs', 'false')
config.set('execution', 'crashfile_format', 'txt')
from nipype import (Node, MapNode, Function, DataGrabber,
                    Workflow, IdentityInterface)
from nipype.interfaces.io import SelectFiles, DataSink

''' define data '''
t_r = 1
vox_size = 1.6

# NEED TO EXTRACT CONTROL ROIs AND PLACE IN REFERENCE DIRECTORY
atlas = 'control_rois'
atlas_roi_files = sorted(glob('/om2/user/ksitek/reference/%s/aparc+aseg-*.nii.gz'%atlas))
#atlas_roi_files = sorted(glob('/om2/user/ksitek/reference/%s/*.nii.gz'%atlas))

#atlas = 'sub-invivo_MNI_rois_like_MNI152_T1_1mm_brain_1p6'
#atlas_roi_files = glob('/om2/user/ksitek/reference/mni_icbm152_nlin_asym_09b/' \
#                       'aud-atlas/atlas_correct-space/' \
#                       'rois_%s/*roi*.nii.gz'%atlas)
# atlas_roi_files = glob('/om2/user/ksitek/reference/mni_icbm152_nlin_asym_09b/' \
#                        'aud-atlas/derivatives/atlas_rois_hcp-1p6/roi*.nii.gz')

''' set smoothing inputs '''
fwhm = vox_size*2

''' define directories '''
analysis = 'rs_corr_fwhm-%.01f_%s_audcort'%(fwhm, atlas)
work_dir = os.path.join('/om2/scratch/', 'Fri', 'ksitek/hcp/', analysis)
project_dir = os.path.abspath('/om2/user/ksitek/hcp_7t/')
out_dir = os.path.join(project_dir, 'derivatives', analysis)

''' define subjects '''
sub_list = [os.path.basename(x) for x in sorted(glob(project_dir+'/data/*'))]

''' set up nodes '''
# set up iterables
infosource = Node(IdentityInterface(fields=['subject_id']),#, 'acq_dir', 'fwhm']),
                                    name='infosource')
infosource.iterables = [('subject_id', sub_list)]#,
                        #('acq_dir', acq_dir_list),
                        #('fwhm', fwhm_list)]

# Create DataGrabber node
dg = Node(DataGrabber(infields=['subject_id'],#, 'acq_dir'],
                      outfields=['anat', 'atlas', 'func']),
          name='datagrabber')

# Location of the dataset folder
dg.inputs.base_directory = os.path.join(project_dir, 'data')

# Necessary default parameters
dg.inputs.template = '*'
dg.inputs.sort_filelist = True

dg.inputs.template_args = {'anat': [['subject_id']],
                           'atlas' : [['subject_id']],
                           'func': [['subject_id']]}
dg.inputs.field_template = {'anat': '%s/MNINonLinear/T1w_restore.1.60.nii.gz',
                            'atlas': '%s/MNINonLinear/ROIs/Atlas_ROIs.1.60.nii.gz',
                            'func': '%s/MNINonLinear/Results/rfMRI_REST*_7T_*/'\
                                    'rfMRI_REST*_7T_*.nii.gz'}

''' create subject-specific mask '''
def make_mask(atlas):
    import nibabel as nib
    import numpy as np
    import os
    #subcort_labels = [16, 28, 60, 10, 49] # brainstem, L/R VentralDC, L/R thalamus

    #roi_labels = [1030, 1034, 2030, 2034] # L/R aud ctx
    roi_labels = [1021, 1028, 2021, 2028] # ctx-lh-pericalcarine, ctx-lh-superiorfrontal, ctx-rh-pericalcarine, ctx-rh-superiorfrontal
    atlas_img = nib.load(atlas)
    atlas_data = atlas_img.get_data()
    affine = atlas_img.affine

    mask_data = np.isin(atlas_data, roi_labels).astype('uint8')
    mask_img = nib.Nifti1Image(mask_data, affine=affine)
    mask_filename = os.path.abspath('roi-control_mask.nii.gz')
    nib.save(mask_img, mask_filename)

    return mask_filename
masker = Node(Function(input_names=['atlas'],
                       output_names=['mask_filename'],
                       function=make_mask), name='masker')


''' extract timeseries from auditory label regions '''
def extract_mean_ts(functional_run, atlas_masks, fwhm):
    import nibabel as nib
    import numpy as np
    from nilearn.input_data import NiftiMasker
    import os

    func_img = nib.load(functional_run)
    func_data = func_img.get_fdata()

    mean_ts = []
    for ax, atlas_mask in enumerate(sorted(atlas_masks)):
        mask_img = nib.load(atlas_mask)
        region_masker = NiftiMasker(mask_img=mask_img, smoothing_fwhm=fwhm,
                                     mask_strategy='epi')
        func_data_2D = region_masker.fit_transform(func_img)

        # compute mean timeseries
        single_mean_ts = func_data_2D.mean(axis=1)

        # save mean time series to txt file
        single_mean_ts_file = os.path.abspath('roi-%02d_mean_ts.txt'%ax)
        np.savetxt(single_mean_ts_file, single_mean_ts, fmt='%f')

        mean_ts.append(single_mean_ts_file)
    return mean_ts
# create node that takes single run and all masks
# (can't do different-length iterfields with MapNode)
mean_ts_extracter = MapNode(Function(input_names=['functional_run',
                                                  'atlas_masks',
                                                  'fwhm'],
                                     output_names=['mean_ts'],
                                     function=extract_mean_ts),
                            name='mean_ts_extracter',
                            iterfield=['functional_run'])
mean_ts_extracter.inputs.fwhm = fwhm
mean_ts_extracter.inputs.atlas_masks = atlas_roi_files


''' datasink '''
ds = Node(DataSink(parameterization=True), name='sinker')
ds.inputs.base_directory = out_dir
ds.plugin_args = {'overwrite': True}

''' create workflow '''
wf = Workflow(name='rs_atlas')
wf.config['execution']['crashfile_format'] = 'txt'

wf.connect(infosource, 'subject_id', dg, 'subject_id')
wf.connect(dg, 'atlas', masker, 'atlas')
wf.connect(dg, 'func', mean_ts_extracter, 'functional_run')

wf.connect(infosource, 'subject_id', ds, 'container')
wf.connect(masker, 'mask_filename', ds, 'subcortical_mask')

wf.connect(mean_ts_extracter, 'mean_ts', ds, 'seed_timeseries')

wf.base_dir = work_dir

wf.run(plugin='MultiProc', plugin_args={'n_procs' : 8})
```
